# The telomere-to-telomere genome and lifestyle transcriptome profiling of *Discula destructiva* Redlin provide modern molecular and genomic context to a historical epidemic

**DOI:** 10.1101/2025.08.04.668385

**Authors:** I. Shade Niece, Jonathan E. Beever, Sonia J. Moisá, Robert N. Trigiano, Kimberly D. Gwinn, William E. Klingeman, Margaret E. Staton, Marcin Nowicki

## Abstract

Fungal pathogens have dramatically altered forests worldwide, yet the mechanisms underlying their virulence remain poorly understood. From the 1970s to the early 2000s, dogwood anthracnose, caused by *Discula destructiva* Redlin, devastated flowering and Pacific dogwoods (*Cornus florida* L. and *C. nuttallii* Aud., respectively). Despite the impacts of *D. destructiva* and other phytopathogens on forest ecosystems, genomic resources remain limited, hindering efforts to understand pathogenicity. The goal of this study was to evaluate the historical *D. destructiva* epidemic through a modern genomics lens by uncovering virulence-associated genes that likely contributed to its rapid spread across native dogwoods. We therefore utilized PacBio HiFi and Proximo Hi-C sequencing to assemble the first telomere-to-telomere, chromosome-scale genome for *D. destructiva* isolate AS111. The resulting 46.655 Mb assembly comprised eight chromosomes with an overall BUSCO completeness of 97.64%. We also identified 10,373 predicted gene models with an overall BUSCO completeness of 96.45%. To investigate gene expression across distinct life cycle phases, reproductive (sporulating) and vegetative (nonsporulating), we conducted RNA sequencing and identified 240 differentially expressed genes (*padj* < 0.05). GO enrichment revealed 162 upregulated genes during sporulation linked to plant cell wall degradation and sugar metabolism, whereas 78 downregulated genes were linked to electron carrier activity and redox balance. Among these 240 genes, 117 genes had predicted protein sequences that were also identified as a candidate virulence factor, including signal peptides, carbohydrate-active enzymes (CAZymes), and effectors, highlighting the role of sporulation-associated gene expression in *D. destructiva* virulence. Together, these findings suggest that the reproductive phase primes *D. destructiva* for host invasion and ecological persistence, which may influence its success as a forest pathogen.

**Author Summary:** Forest pathogens threaten ecosystems worldwide and cause extensive ecological and economic damage. Since the 1970s, native dogwood populations in North America have been devastated by dogwood anthracnose, caused by the exotic fungal pathogen *Discula destructiva*. *Discula destructiva* is just one of many destructive fungal pathogens in the order Diaporthales, which includes other noteworthy pathogens that cause chestnut blight (*Cryphonectria parasitica*) and butternut canker (*Ophiognomonia clavigignenti-juglandacearum*). Despite the widespread impact of fungal diseases, little is known about the genetic factors that drive their spread and severity. To help address this gap, we generated the first telomere-to-telomere, chromosome-scale genome assembly of *D. destructiva* and analyzed gene expression across distinct life cycle phases: reproductive (sporulating) and vegetative (nonsporulating) growth. Our findings revealed key sporulation-associated genes and virulence factors that may facilitate host infection and underscore the importance of sporulation in the pathogenicity of *D. destructiva*. These insights improve our understanding of the mechanisms that drive disease development, influence how fungal pathogens such as *D. destructiva* establish and persist in forest ecosystems, and provide a foundation for future comparative genomics among other devastating pathogens in Diaporthales.

## Introduction

Fungi in the order Diaporthales (Sordariomycetes, Ascomycota) have had substantial impacts on natural and agricultural ecosystems globally [1]. This lineage includes phytopathogens (plant pathogens) specialized in colonizing leaves, stems, and bark (plant tissues critical for growth and survival) and is associated with symptoms ranging from superficial blemishes to rapid host decline [2,3]. Some of the most notable Diaporthalean forest pathogens in North America include the causal agents of chestnut blight (*Cryphonectria parasitica* Murill Barr), butternut canker (*Ophiognomonia clavigignenti-juglandacearum* Broders & Bolan), and dogwood anthracnose (*Discula destructiva* Redlin) [2–4]. *Castanea dentata* Borkh. (American chestnut) and *Juglans cinerea* L. (butternut) have been driven to functional extinction and critical endangerment by chestnut blight and butternut canker, respectively [5]. Despite an estimated loss of 4.6 billion trees to the initial dogwood anthracnose epidemic, *Cornus florida* L. (flowering dogwood) has shown notable persistence, likely due to a combination of host–pathogen and host–environment interactions [6,7]. The genus *Cornus* (dogwood) includes about 12 species in North America, where two species are most susceptible to *D. destructiva*: *C. florida* L. (flowering dogwood) and *C. nuttallii* Aud. (Pacific dogwood) [8]. Both *C. florida* and *C. nuttallii* lack widespread resistance to *D. destructiva*, which resulted in one of the largest forest epidemics of the 20th century in North America [4,9–11]. *Cornus florida* forms second-growth hardwood stands in eastern North America that provide important ecosystem services, including wildlife nourishment and calcium cycling, and serve as a key understory component of old-growth forests [12–15]. *Cornus nuttallii* occupies a similar understory role in riparian forests of the Pacific Northwest [16]. Additionally, *C. kousa* Hance (kousa dogwood), a species native to eastern Asia, has gained popularity in North America for its ornamental appeal, including within urban forests and managed landscapes [8,9,17]. Numerous species and several hundred cultivars of *Cornus* are planted as woody ornamentals, which generate an annual revenue of nearly $30 million USD in deciduous flowering tree sales in the United States of America (USA) [17–21].

Symptoms of dogwood anthracnose range from brown, necrotic lesions on bracts and leaves to the formation of cankers that coalesce and girdle the trees, ultimately leading to canopy dieback and mortality [4,9,22–24]. Mortality may occur in as little as two to three years after initial infection, with saplings and understory trees being the most susceptible to infection [23–26]. The first case of *D. destructiva* was reported for *C. nuttallii* in Washington in 1976, though its identity remained uncertain until its formal description in 1991 [4,23,27,28]. Shortly after the initial 1976 report on the western coast of the USA, *D. destructiva* was documented in *C. florida* on the eastern coast in New York and Connecticut in 1977 [4,23,24,27,28]. An additional fungus commonly co-isolated with *D. destructiva* was later shown to induce similar disease symptoms in *Cornus*, initially classified as *Discula* Type II and later reclassified as *Juglanconis japonica* [9,29]. *Juglanconis japonica* is distinguished by differences in morphology, lack of polyphenol oxidase production (a key enzyme produced during *D. destructiva* infection involved in lignin degradation), and ITS sequence divergence [23,29–33].

*Discula destructiva* exists strictly as a hemibiotrophic, haploid anamorph with no observed sexual stage *in vitro* or *in vivo* and relies solely on conidia (asexual spores) for survival and reproduction [4,34]. In the initial biotrophic phase, *D. destructiva* relies on living host tissue for nutrient acquisition. However, late-stage infection is characterized by a life cycle phase transition to necrotrophy, during which enzymes and toxins cause tissue death [32,35–37]. This flexible life cycle may have enabled the rapid spread of *D. destructiva* across North America, which previous work suggests was facilitated by conidia dispersal via migratory birds, animals, and–markedly–the horticultural trade [26,38,39]. As a result of this rapid dispersal across a wide geographic range and a lack of natural resistance in affected *Cornus* species, an estimated 3 billion *C. florida* succumbed to *D. destructiva* in the Appalachian ecoregion alone between 1977 and 2010 [7,25]. Ultimately, the *D. destructiva* epidemic left disjunct populations of *C. nuttallii* critically imperiled in Idaho and fragmented populations of *C. florida* critically imperiled in southeastern Canada and parts of the eastern USA [40–42].

The prevalence of *D. destructiva* and dogwood anthracnose has diminished in cultivated landscapes in recent years, due in part to the widespread use of tolerant germplasm and cultural control practices such as pruning infected tissues [43,44]. To date, only one truly resistant *C. florida* cultivar, ‘Appalachian Spring’, has been identified, which was originally selected from a survivor tree in Catoctin Mountain Park, Maryland [22]. Similarly, disease pressure in natural *Cornus* populations has diminished, likely due to the extirpation of highly susceptible hosts [43,45]. Although *D. destructiva* currently poses a less immediate threat, its epidemic history provides an opportunity to apply modern genomic techniques to inform future breeding and management strategies for dogwood anthracnose, as well as develop improved control methods for related *Discula* species that infect various ecologically and economically important trees such as *Fraxinus*, *Acer*, *Fagus*, *Platanus*, and *Quercus*. This wide range of plant hosts susceptible to *Discula* species highlights the broader relevance of understanding pathogenicity within the genus. As the global movement of pathogens continues to accelerate due to climate change and trade, there is an urgent need for proactive approaches to facilitate forest pathogen surveillance and mitigation [46]. Understanding genetic diversity and its influence on virulence and host-pathogen interactions in known epidemic systems, such as *D. destructiva*, provides an opportunity to develop future predictive frameworks for identifying and responding to potential threats. A previous population genetics study using microsatellites suggested four geographically and temporally distinct genetic clusters in *D. destructiva* across the USA and implicated *C. kousa* as the source of multiple introductions [9–11,47–49]. Currently, only five chromosome-scale genome assemblies in the order Diaporthales have been published on the National Center for Biotechnology Information (NCBI) (Table S1), and none existed for *Discula* prior to this study [50]. As long-read sequencing continues to become more accessible, developing high-quality genome resources is a valuable and increasingly tractable first step toward profiling the invasion potential of pathogens. The objective of this research was to resolve this knowledge gap for a forest pathogen with known consequences and epidemic potential by assembling the first annotated, long-read reference genome for *D. destructiva*. Synergistically, RNA sequencing (RNAseq; Table S2) was conducted to identify candidate genes of importance during sporulation, to provide insights into both mechanisms of sporulation and its contribution towards pathogenicity. Additionally, we leveraged these resources to identify candidate virulence factors (signal peptides, carbohydrate-active enzymes (CAZymes), and effectors) that may compromise host defenses and support fungal growth. Together, these analyses provide genomic insights into the mechanisms driving *D. destructiva*’s historical spread and support future comparative genomic studies across *Discula* and Diaporthales more broadly.

## Materials & Methods

### Culture conditions of fungal isolates & extractions

Two *D. destructiva* isolates (AS111 and WAP31) and one *J. japonica* isolate (VA17B), all initially collected from infected flowering and Pacific dogwoods from 1989 to 2000 (Table 1), were utilized throughout this study as resources for high molecular weight (HMW) DNA and RNA extractions. The isolates of *D. destructiva* represent separate genetic clusters that originated from *C. florida* (AS111) and *C. nuttallii* (WAP31) (Table 1) [9]. The *J. japonica* isolate was also chosen as a comparator due to its original description as *Discula* Type II that presents unique metabolic characteristics [9,24,32]. Axenic stock cultures of each isolate were maintained on solid half-strength potato dextrose agar (Thermo Scientific; Hampton, NH, USA) amended with chlortetracycline hydrochloride and streptomycin sulfate (PDA^++^). Stock cultures were incubated at room temperature (ca. 21 °C) under constant light for up to 14 days prior to being transferred to growth media for subsequent HMW DNA and RNA extractions.

**Table 1.** Metadata for the Discula destructiva and Juglanconis japonica isolates.

| **Species<sup>a</sup>** | **Isolate** | **Location/Year<sup>b</sup>** | **Collection<sup>c</sup>** | **Host Species** | **State Isolated** |
| --- | --- | --- | --- | --- | --- |
| *Discula destructiva* | AS111 | S/2000 | UTK | *Cornus florida* | Tennessee |
| *Discula destructiva* | WAP31 | W/1989 | RU | *Cornus nuttallii* | Washington |
| *Juglanconis japonica* | VA17B | NA/1991 | UTK | *Cornus florida* | Virginia |

<sup>a</sup> Original classification of *Juglanconis japonica* as *Discula* Type II.
<sup>b</sup> Geographic locations where isolates were originally collected with the collection year: S = South USA; W = West USA; NA=not applicable. Locations denoted per Mantooth et al., 2017 [9].
<sup>c</sup> Original source of fungal isolates: UTK = University of Tennessee (Knoxville, TN, USA); RU = Rutgers University (New Brunswick, NJ, USA).

For HMW DNA extractions, 5 mm^2^ mycelial plugs were cut from margins of actively growing stock cultures of *D. destructiva* isolate AS111, then individually transferred to sterilized cellophane disks (BIO-RAD Laboratories; Hercules, CA, USA) overlaid on solid half-strength PDA^++^ [50]. Cultures were incubated at room temperature (ca. 21 °C) under constant light for up to 14 days prior to extractions. After incubation, mycelium was scraped off the cellophane and submerged into liquid nitrogen. A sterilized mortar and pestle was used to homogenize frozen mycelium into a fine powder. HMW DNA was extracted using a modified protocol of the Qiagen MagAttract HMW DNA Kit (Qiagen; Valencia, CA, USA), and excess polysaccharides were removed from the extracted genomic DNA. HMW DNA extraction and polysaccharide precipitation protocols are available in supplemental materials (File S1 and File S2, respectively). The quality and quantity of the purified genomic DNA were assessed on a FEMTO Pulse System (Agilent; Santa Clara, CA, USA), and DNA was stored at -20 °C until library preparation and sequencing.

For RNA extractions, 5 mm^2^ mycelial plugs were cut from margins of actively growing stock cultures of two *D. destructiva* isolates (AS111 and WAP31) and one *J. japonica* isolate (VA17B), then individually transferred to sterilized cellophane disks overlaid on two different media types: solid half-strength PDA^++^ and solid water agar supplemented with finely ground *Quercus alba* L. (white oak) leaf tissue [51]. These growth media were selected as the non-sporulating (strictly of vegetative, mycelial growth) and sporulating (mycelia and abundant acervuli present) media, respectively, used to enable detection of differential gene expression between distinct life cycle phases (Fig S1). Cultures were incubated at room temperature (ca. 21 °C) under constant light for up to 14 days prior to RNA extractions. After incubation, RNA was extracted using a custom TRIzol™ Reagent (Invitrogen; Waltham, MA, USA) protocol (File S3).

Total RNA quality and quantity were assessed using a TapeStation High Sensitivity RNA ScreenTape (Agilent) and stored at -20 °C until library preparation and sequencing.

### Library preparation & sequencing

Prior to PacBio High Fidelity (HiFi) long-read sequencing (University of Washington Northwest Genomics Center; Seattle, WA, USA), purified genomic DNA from *D. destructiva* isolate AS111 was fragmented into 14 kilobase (kb) fragments using the PippenHT instrument. HiFi libraries were constructed using the PacBio SMRTbell Prep Kit 3.0, then sequenced on the PacBio Revio system. To detect genome-wide chromatin interactions within nuclei, raw mycelial tissue obtained from stock cultures of *D. destructiva* isolate AS111 was used as input for Hi-C sequencing. The Proximo high-throughput Chromosome Conformation Capture (Hi-C) Fungal Kit Protocol (Phase Genomics; Seattle, WA, USA) was used to extract DNA and prepare libraries for paired-end sequencing (2 × 150 bp) on an Illumina platform. The following combination of restriction enzymes were used for Hi-C library preparation: DpnII (^GATC), Ddel (C^TNAG), MseI (T^TAA), and Hinfl (G^ANTC).

The RNAseq experimental design consisted of three isolates, each represented by three biological replicates, exposed to two treatment conditions (non-sporulating and sporulating), resulting in a total of 18 samples. To identify differentially expressed genes between treatments, total RNA was processed for library preparation using the SMART-Seq mRNA LP (with UMIs) Kit (Takara Bio Inc; Kusatsu, Shiga, Japan) and enriched for mRNA via poly(A) selection.

Enriched mRNA was reverse transcribed into 18 complementary DNA libraries, which were sequenced (2 × 150 bp) on an Illumina NovaSeq platform (University of Tennessee Genomics Core; Knoxville, TN, USA).

### Telomere-to-telomere (T2T), chromosome-scale genome assembly

FastQC v0.11.7 [52] was used to perform quality control checks on the raw Hi-C sequencing data. To reduce computational resources and data redundancy during genome assembly, the raw HiFi sequencing data was downsampled from approximately 1,900× coverage to 100× coverage using seqtk v1.2-r94 [53]. Using the downsampled 100× HiFi reads and raw Hi-C reads, Hifiasm v0.19.8 [54] was used to assemble a draft genome assembly with the following parameters: -t 32, --hg-size 49m, --n-hap 1, --n-weight 10, --f-perturb 100000, --n- perturb 0.25, --h1 $hic_r1, --h2 $hic_r2. Next, potential contamination was identified by first using the NCBI Foreign Contamination Screen (FCS) tool suite. Specifically, FCS-GX v0.5.0 [55] and FCS-ADAPTOR v0.5.0 [50] were used with default parameters to identify and remove (if present in the assembly) any contaminant sequences derived from other organisms or sequencing adaptors. To filter out mitochondrial sequences from the nuclear genome assembly, the mitochondrial genome was first *de novo* assembled using Oatk v1.0 [56] with default parameters. To assess the assembly completeness and gene predictions for the mitochondrial genome, BBMap v39.06 [57] and GeSeq v2.03 [58] were used with default parameters. GeSeq annotations were based on the closest related Diaporthales reference species: *Chrysoporthe austroafricana* (NC_030522.1), *C. cubensis* (NC_030524.1), and *C. deuterocubensis* (NC_030523.1). OGDRAW v1.3.1 [59] was used to visualize the annotated mitochondrial genome. Mitochondrial sequences present in the nuclear assembly were then filtered out by running a custom Python script (File S4) that identified and removed aligned regions with more than 90% alignment length. Hi-C scaffolding was then performed on the filtered nuclear genome assembly using YaHS v1.2a.2-2 with default parameters and the following restriction enzyme motifs specified: GATC,CTNAG,TTAA,GANTC [60]. Scaffolds were manually corrected and visualized using JBAT v2.15 [61]. Assembly gaps were resolved by aligning the raw HiFi reads to the scaffolded assembly using TGS-GapCloser v1.1.1 default parameters [62]. To determine whether the chromosome-level scaffolds represented a T2T assembly, tidk 0.2.65 was used with the “search” function and these telomeric motifs specified: 5’-(TTAGGG/CCCTAA)_n_-3’ [63]. Final genome assembly statistics were generated using BBMap v39.06 [57]. Genome assembly completeness was assessed using the Benchmarking Universal Single-Copy Orthologs (BUSCO) v5.5.0 [64] sordariomycetes_odb10 database and ascomycota_odb10 database. Likewise, the compleasm v0.2.6 [65] sordariomycetes_odb10 database and ascomycota_odb10 database were used to provide an additional completeness estimate. Finally, Merqury v1.3-2 [66] was used to assess genome assembly completeness and quality using a k-mer-based, reference-free approach.

### Genome annotation

The newly constructed *D. destructiva* AS111 reference genome was annotated with the following steps. Repetitive elements were identified and classified using RepeatModeler v2.0.5 with default parameters and the -LTRStruct flag specified [67]. These repeat elements were then soft masked using RepeatMasker v4.1.6 with default parameters and these optional parameters: - e rmblast, -pa 4, -nolow, -xsmall, -gff. [68]. Raw RNAseq read-pairs were aligned to the *D. destructiva* AS111 reference genome using STAR v2.7.11b with default parameters and these optional parameters: --outSAMtype BAM SortedByCoordinate and --outSAMstrandField intronMotif. [69]. Structural annotations were performed using BRAKER v3.0.8 with default parameters and the --fungus flag specified [70]. The BUSCO sordariomycetes_odb10 database and ascomycota_odb10 database were used to assess the completeness of the predicted proteome output from BRAKER. Finally, functional gene predictions were performed using EnTAP v1.0.0 with default parameters [71].

### Differential gene expression analysis

FastQC v0.11.7 [52] was used to perform quality control checks on raw RNAseq read-pairs. Read trimming was performed with BBDuk v39.06 [57] to remove any Illumina adaptor contamination and any sequences with a Phred quality score lower than 30. Default parameters were used for BBDuk with these optional parameters specified: trimpolya=10, trimpolyg=10. Residual adaptor contamination was subjected to further alignment filtering and exclusion using STAR. Trimmed RNAseq read-pairs were then aligned to the *D. destructiva* AS111 reference genome using STAR with default parameters and the optional -- outSAMtypeBAMSortedByCoordinate flag specified. The number of RNAseq reads that mapped to unique genomic features was calculated using FeatureCounts v2.0.6 with these parameters: -p, -s 2, -T 30, -t gene, -g ID, -F GTF, -a dd.gff3, -o combined.counts.txt, *bam [72]. To perform differential gene expression analysis between sporulating and non-sporulating treatments, the gene count matrix produced from FeatureCounts was used to run the DESeq2 v1.42.1 [73] pipeline in RStudio v2024.12.0.467 (File S5) [74]. To identify genes with the most expression changes between treatments, a confidence level of 95% (α = 0.05) was applied, allowing for the determination of significantly upregulated and downregulated genes (*padj* < 0.05). A stringent filter of *padj* = 1×10^−7^ and *log_2_FC* = 2 was then applied to identify the most significant subset of differentially expressed genes.

### Gene Ontology (GO) enrichment

All significantly differentially expressed genes (*padj* < 0.05) were used as input for GO enrichment analysis in RStudio to identify enriched GO terms among upregulated and downregulated genes (File S6). Additionally, the most significant subset of differentially expressed genes (*P value* = 1×10^−7^ and *log_2_FC* = 2 ) were analyzed to identify associated Kyoto Encyclopedia of Genes and Genomes (KEGG) Ontology (KO) annotations predicted during functional annotation [75]. Genes that lacked annotated KO annotations were further analyzed using these additional protein similarly search resources: GhostKOALA v3.1 [76], BlastKOALA v3.2 [76], and InterProScan [77], and local NCBI BLASTp [50].

### Identification of candidate virulence factors

To identify candidate virulence factors from the predicted proteome of the *D. destructiva* AS111 reference genome, SignalP v5.0 [78], dbCAN v4.1.4 [79], and EffectorP v3.0 [80] were independently run to identify candidate signal peptides, CAZymes, and effectors, respectively. The proteome was utilized to ensure that divergent gene model protein predictions were retained for a comprehensive prediction of these virulence factors. Default parameters were used for all three tools, with these optional SignalP parameters -format long, -gff3, -mature and the optional EffectorP -f flag specified. We used a custom R script (File S7) to determine if the predicted protein sequences of any differentially expressed sporulation genes (*padj* < 0.05) were also predicted as candidate virulence factors.

## Results

### Generation of a T2T, chromosome-scale genome assembly for Discula destructiva

To accomplish a T2T, chromosome-scale assembly of the *D. destructiva* genome, we generated approximately 5.8 million raw HiFi reads (93.51 GB) from SMRTbell libraries prepared from isolate AS111 (Table S3). Raw HiFi genome sequencing was estimated to be 1,893× coverage (Table S3). Hi-C libraries for *D. destructiva* isolate AS111 generated approximately 50.6 million raw read-pairs with an average per sequence quality score of Q39 and no adaptor contamination. Hifiasm generated a 46,885,455 bp contig-level genome assembly with ten scaffolds and 10 gaps present using the 100× downsampled HiFi data and raw Hi-C data (Table S4). No contamination sequences or adaptors were identified during the contamination screening using the NCBI FCS tool suite. Oatk generated a 232,813 bp mitochondrial genome assembly that consisted of one circularized contig (Fig 1A). Gene prediction using GeSeq revealed six protein coding gene families in the mitochondrial genome assembly (Fig 1A). After mitochondrial sequence removal from the nuclear assembly, scaffolding of the nuclear assembly using YaHS and manual curation in JBAT resulted in eight chromosome-level scaffolds, with 100% of the contigs placed within these eight chromosome-level scaffolds and eight gaps present (Table S4 and Fig 1B). TGS-GapCloser produced a final 46,654,619 bp chromosome-level genome assembly with zero gaps (Table 2 and Fig 1C). After running tidk to determine if the chromosome-scale scaffolds represented a T2T assembly, telomeric motifs 5’-(TTAGGG/CCCTAA)_n_-3 were identified on both ends of all eight chromosome-level scaffolds (Fig S2). There were an average of 32.6 and 32.1 forward- and reverse telomeric motifs 5’- (TTAGGG/CCCTAA)_n_-3, respectively. Assessment of genome completeness with the BUSCO sordariomycetes_odb10 and ascomycota_odb10 databases on the final T2T, chromosome-level genome assembly indicated a high completeness score of 95.60% and 97.40%, respectively (Table 2). Assessment of genome completeness with the compleasm Sordariomycetes and Ascomycota lineage datasets (sordariomycetes_odb10 and ascomycota_odb10, respectively) also indicated a high completeness score of 97.30% and 99.47%, respectively. Assessment of genome completeness with Merqury indicated a high completeness score of 98.45%, with an estimated assembly consensus quality value of Q79 and an error rate of 1.26 × 10^-8^.

**Fig 1.**
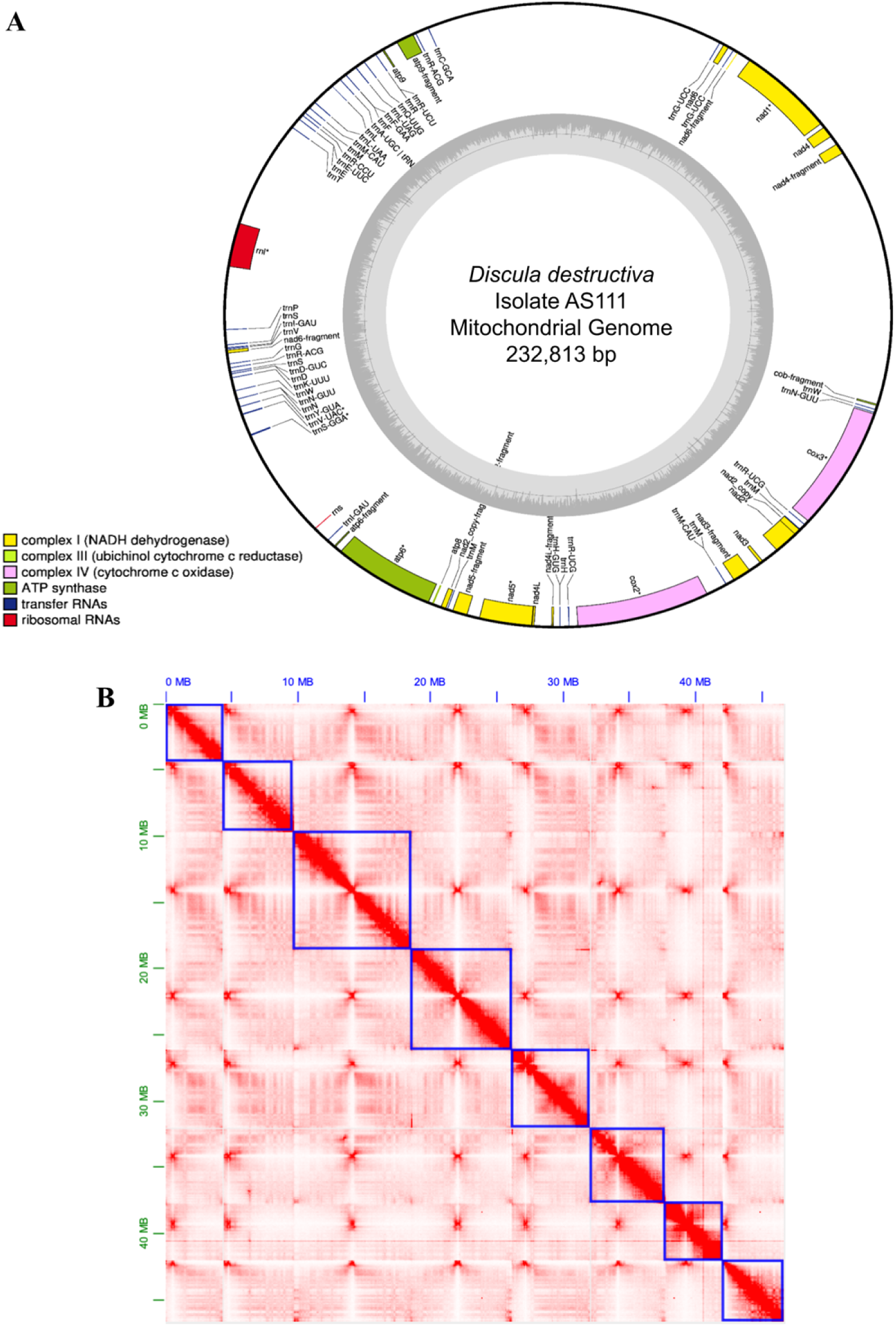
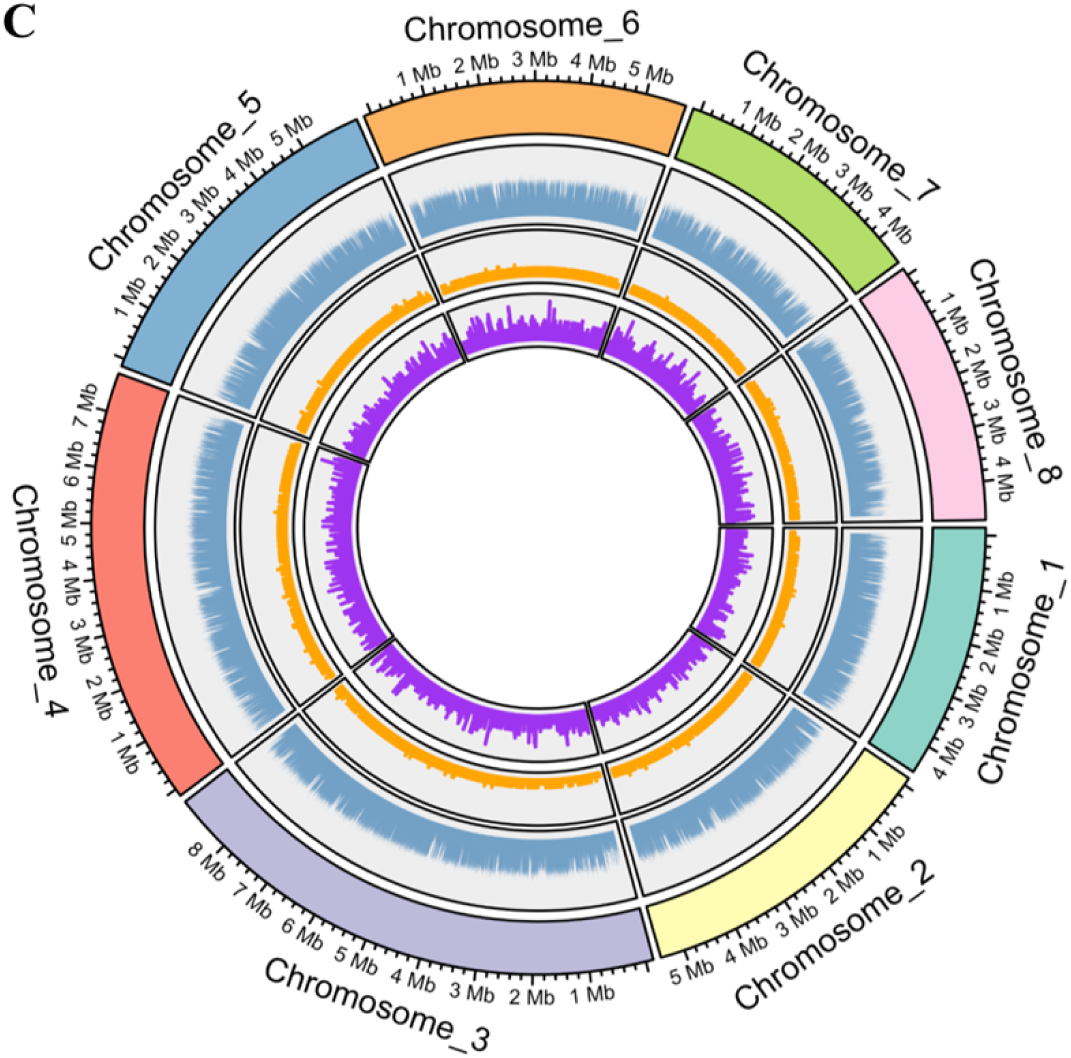
Overview of the mitochondrial and nuclear genome assemblies of *Discula destructiva* isolate AS111. **A.** The 232,813 bp circularized mitochondrial genome assembly generated by Oatk v1.0 that consists of one, circularized contig. Genome annotations generated using GeSeq v2.02 and visualized using OGDRAW v1.3.1. Annotations of predicted gene families (lower left corner) are indicated by the colored bars, and GC content is represented by the inner, grey circle. **B.** Hi-C contact map generated using Juicebox Assembly Tools (JBAT) v2.15 displaying the genomic interactions across the final telomere-to-telomere, chromosome-scale nuclear genome assembly of 46.655 megabases (Mb). The blue boxes outline the eight chromosome-level scaffolds in the assembly. Both the x- and y-axes are labeled with genomic coordinates in Mb. **C.** Circlize v0.4.16 plot of the final T2T, chromosome-scale nuclear genome assembly. From outer to inner rings, we depict: chromosome numbers with chromosome size indicated by Mb tick marks, a histogram of percentage of GC content in 1 kilobase (kb) windows, a line plot of gene density in 1 kb windows, and a bar plot of transposable element density in 1 kb windows.

**Table 2.**
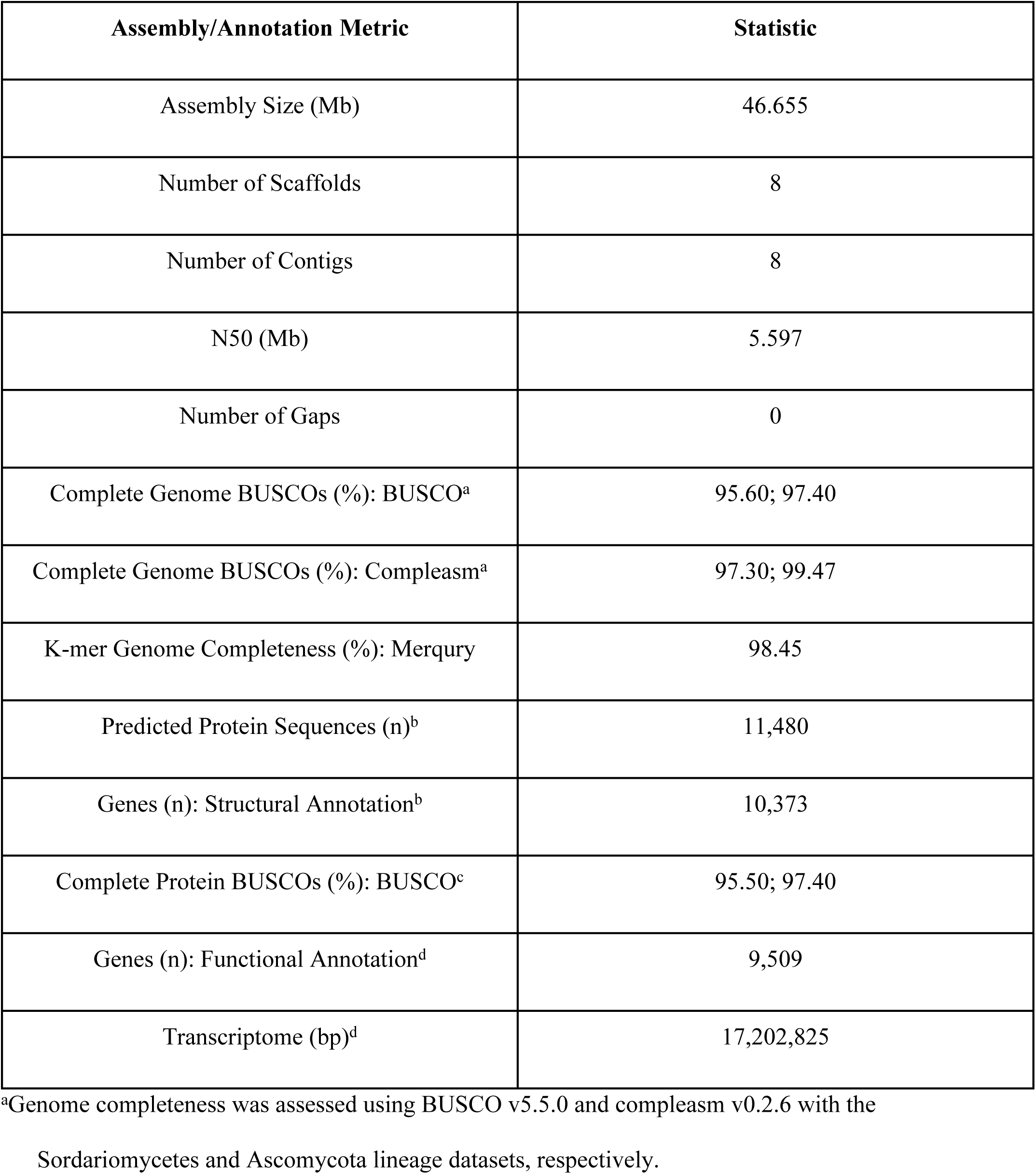
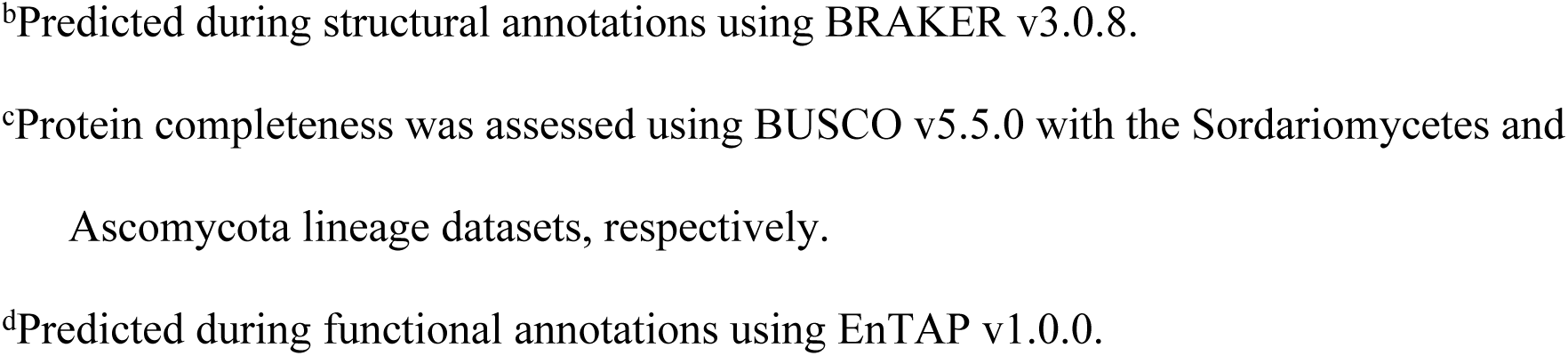
Genome assembly and annotation statistics for the final telomere-to-telomere, chromosome-scale nuclear genome assembly of *Discula destructiva* isolate AS111.

### Structural and functional genome annotation

RepeatModeler predicted a total of 135 repeat elements, of which 63 (46.7%) were classified into known families, whereas 72 (53.3%) remained unclassified (Table S5). A total of 7,212,922 bp (15.38%) of the 46.655 Mb genome assembly were masked as repeat elements by RepeatMasker. The primary repeat elements masked in the genome assembly were retroelements, specifically LTR elements Ty1/Copia and Gypsy/DIRS1 (Table S5). A total of 11,480 protein sequences and 10,373 genes were predicted during structural annotations with BRAKER (Table 2). Assessment of gene completeness from structural annotation with the BUSCO sordariomycetes_odb10 and ascomycota_odb10 databases indicated a high completeness score of 95.50% and 97.40%, respectively (Table 2). EnTAP functionally annotated 9,509 genes (Table 2), resulting in 91.67% of genes being retained from structural annotation to functional annotation. EnTAP identified 8,330 unique sequences with at least one GO term and 2,725 unique sequences with at least one KEGG pathway assignment. A transcriptome assembly produced from EnTAP was estimated to be 17,202,825 bp (Table 2) with an N50 of 1,770 bp.

### Transcriptomic responses to sporulation

Illumina sequencing of 18 cDNA libraries from *D. destructiva* isolates AS111 and WAP31 and *J. japonica* isolate VA17B generated 94.24 GB of raw RNAseq data, with 1,181,440,126 reads (98.54%) retained after trimming adaptor contamination (Table S6). RNAseq alignment to the *D. destructiva* AS111 reference genome revealed taxon-specific mapping efficiency, with less uniquely mapped reads for *J. japonica* VA17B compared to both *D. destructiva* isolates (Table S7). Differential expression analysis with DESeq2 identified a total of 324 differentially expressed genes (default: *padj* < 0.1) under the sporulation treatment, with 115 upregulated genes and 209 downregulated genes. Based on principal component analysis (PCA) and hierarchical clustering, gene expression profiles segregated greatest by interspecific diversity, with treatment groups clustering together (Fig S3). After filtering for significant differentially expressed genes (*padj* < 0.05), a total of 240 genes were retained, of which 162 genes were upregulated and 78 were downregulated during sporulation (Table S8). To more precisely analyze processes most impacted by sporulation, a stringent filter (*padj* = 1e-7, *log_2_FC* = 2) was applied to the 240 gene set and refined to a subset of the four most upregulated and five most downregulated genes (Fig 2A). A heatmap of the top 40 differentially expressed genes illustrated distinct gene expression patterns, highlighting two major gene clusters with largely inverse expression shifts between treatment groups (Fig 2B). Although the overall expression profiles were similar, some variation of expression intensity could be observed between *D. destructiva* isolates, such as in *g5344* on sporulation media and *g4351* on non-sporulation media (Fig. 2B). Additionally, opposite expression profiles could be observed in some genes between the *D. destructiva* isolates and *J. japonica* isolate. For example, *g9868* and *g9869* were highly downregulated under the sporulation treatment in both *D. destructiva* isolates but upregulated in *J. japonica* isolate VA17B (Fig. 2B).

**Fig 2.**
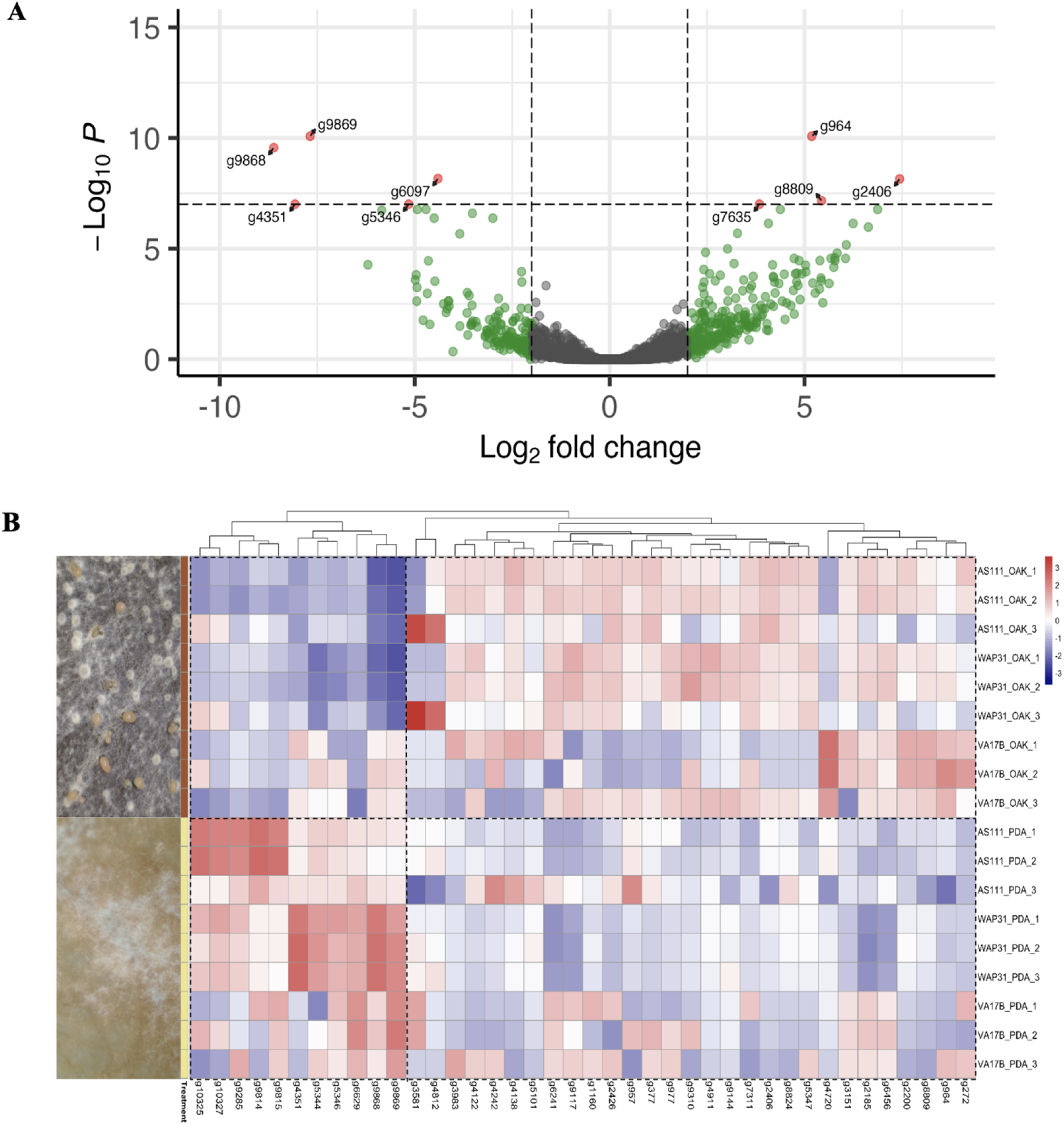
Key sporulation genes and expression patterns in *Discula destructiv*a (AS111, WAP31) and *Juglanconis japonica* (VA17B). **A.** Subset of the nine most differentially expressed genes (*padj* = 1e-7; *log_2_FC* = 2), indicated by red points on the volcano plot. Points to the left indicate the most downregulated genes, whereas points to the right indicate the most upregulated genes during sporulation. Green points indicate genes that meet the *log_2_FC* = 2 threshold but not the *adjusted P value* ≤ 1e-7 threshold, and grey points represent non-significant genes. **B.** Heatmap displaying the expression patterns of the top 40 differentially expressed genes (based on *log_2_FC*) across all species and isolates grown on both sporulation (brown) and non-sporulation (yellow) media. Representative colony images of fungal cultures grown on each medium are shown on the left for reference. Red indicates higher gene expression levels, whereas blue represents lower expression levels. Clustering patterns, indicated via dashed lines, highlight differential gene expression between treatments (top versus bottom dashed boxes by treatment).

### Functional enrichment analysis of sporulation genes

The 162 upregulated sporulation genes (*padj* < 0.05) were enriched for processes involving hydrolase activity, carbohydrate metabolic and catabolic processes, and polysaccharide metabolic and catabolic processes (Fig 3A). The 78 downregulated sporulation genes (*padj* < 0.05) were enriched for processes involving heme binding, tetrapyrrole binding, electron carrier activity, and iron ion binding (Fig 3B). KO annotations associated with the four most upregulated genes (*padj* = 1e-7; *log_2_FC* = 2) were involved in sugar transportation and catalysis of cell wall components (Table 3 and Table S9). KO annotations associated with the five most downregulated genes (*padj* = 1e-7; *log_2_FC* = 2) were involved in the destruction of radicals and ferric-chelate reductase (Table 3). Notably, among these downregulated genes, *g9868* did not have a KO annotation (Table 3). Additional searches using GhostKOALA, BLASTKOALA, and InterProScan found no similarity sequence matches to aid in identifying a functional annotation for *g9868*. A local NCBI BLASTp search identified *g9868* as a hypothetical protein (*F5883DRAFT_551733*, *Diaporthe sp. PMI_573*), whereas the KEGG database BLASTp search returned a similar result for a hypothetical protein (*VDAG_07229*, *Verticillium dahliae*). Neither of the hypothetical proteins had an associated KO number or annotation.

**Fig 3.**
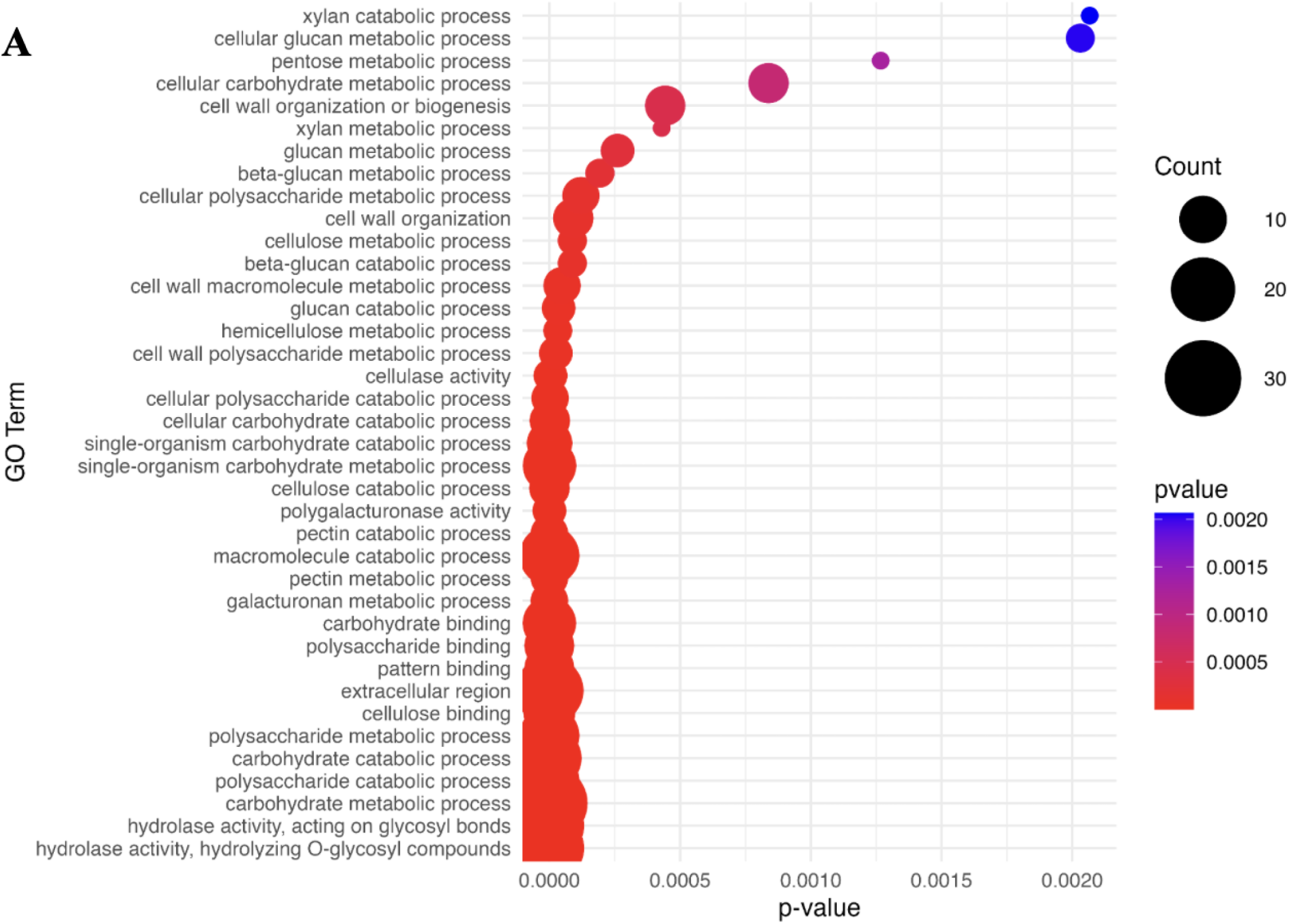
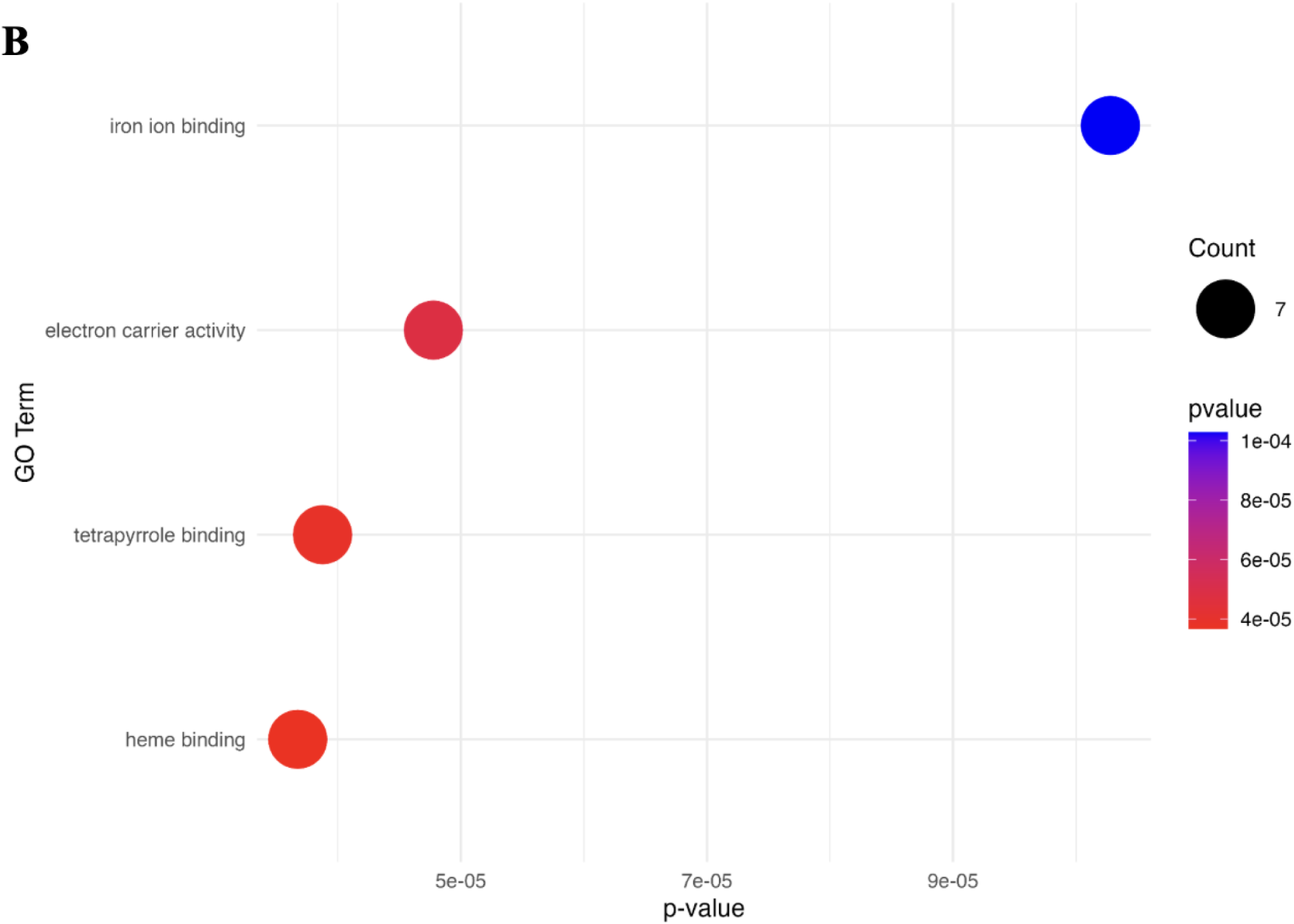
Gene Ontology (GO) enrichment of upregulated and downregulated sporulation genes in *Discula destructiva* and *Juglanconis japonica*. A. Bubble plot displaying the most enriched GO terms among the 162 upregulated sporulation genes (*padj* < 0.05). **B.** Bubble plot displaying the most enriched GO terms among the 78 downregulated sporulation genes (*padj* < 0.05). GO enrichment for both upregulated and downregulated genes was performed using GO terms provided from EnTAP v1.0.0.

**Table 3.** Subset of the most differentially expressed genes and their associated Kyoto Encyclopedia of Genes and Genomes (KEGG) Orthology (KO) annotations.

| Gene ID | KO Annotation | Expression Pattern During Sporulation |
| --- | --- | --- |
| *g9869* | Destroys radicals which are normally produced within cells that are toxic to biological systems (By similarity) | Downregulation |
| *g9868* | N/A | Downregulation |
| *g6097* | Ferric-chelate reductase | Downregulation |
| *g5346* | NAD/NADP octopine nopaline dehydrogenase | Downregulation |
| *g4351* | Ctr copper transporter family | Downregulation |
| *g8809* | Catalyzes endohydrolysis of 1,4-beta-D-glucosidic linkages in xyloglucan with retention of the beta-configuration of the glycosyl residues. Specific for xyloglucan and does not hydrolyze other cell wall components | Upregulation |
| *g7635* | Sugar (and other) transporter | Upregulation |
| *g2406* | Arabinan endo-1,5-α-L-arabinosidase | Upregulation |
| *g964* | Inositol oxygenase | Upregulation |

Subset of the nine most differentially expressed genes ( $p_{adj} = 1e-7$ ; $\log_2FC = 2$ ) and their KO annotations. “N/A” represents a KO annotation that could not be determined from similarity searches using EnTAP v1.0.0, GhostKOALA v3.1, BLASTKOALA v3.2, and InterProScan.

### Secreted proteins, CAZymes, and effectors as candidate virulence genes

Out of the 11,480 predicted protein sequences, SignalP predicted a total of 1,209 signal peptides, which were classified as secretory substrates cleaved by SPase I (Sec/SPI). The remaining 10,271 protein sequences were not classified as signal peptides (Table S10 and Table S11). A total of 665 CAZymes were predicted using dbCAN, with the remaining 10,815 protein sequences lacking a CAZyme prediction (Table S10 and Table S12). These enzymes were assigned to a total of six CAZyme families based on support from at least two annotation tools integrated into dbCAN, following the authors’ recommended criteria. EffectorP predicted a total of 2,763 effectors with the remaining 8,717 protein sequences predicted as non-effectors (Table S10 and Table S13). Among the 240 differentially expressed sporulation genes (*padj* < 0.05), a total of 117 genes were identified to have predicted protein sequences that overlapped with at least one of the three virulence factors considered here (Fig 4 and Table S14). Of those, 25 genes encoded proteins predicted to be all three classes of virulence factors, with 24 genes upregulated and one gene downregulated during sporulation (Fig 4 and Table S14). Notably, *g2406*, *g8809*, *g9868*, and *g9869* from the subset of the nine most differentially expressed sporulation genes (*padj* = 1e-7; *log_2_FC* = 2) (Fig 2A, Table 3) were identified to have overlap as predicted virulence factors.

**Fig 4.**
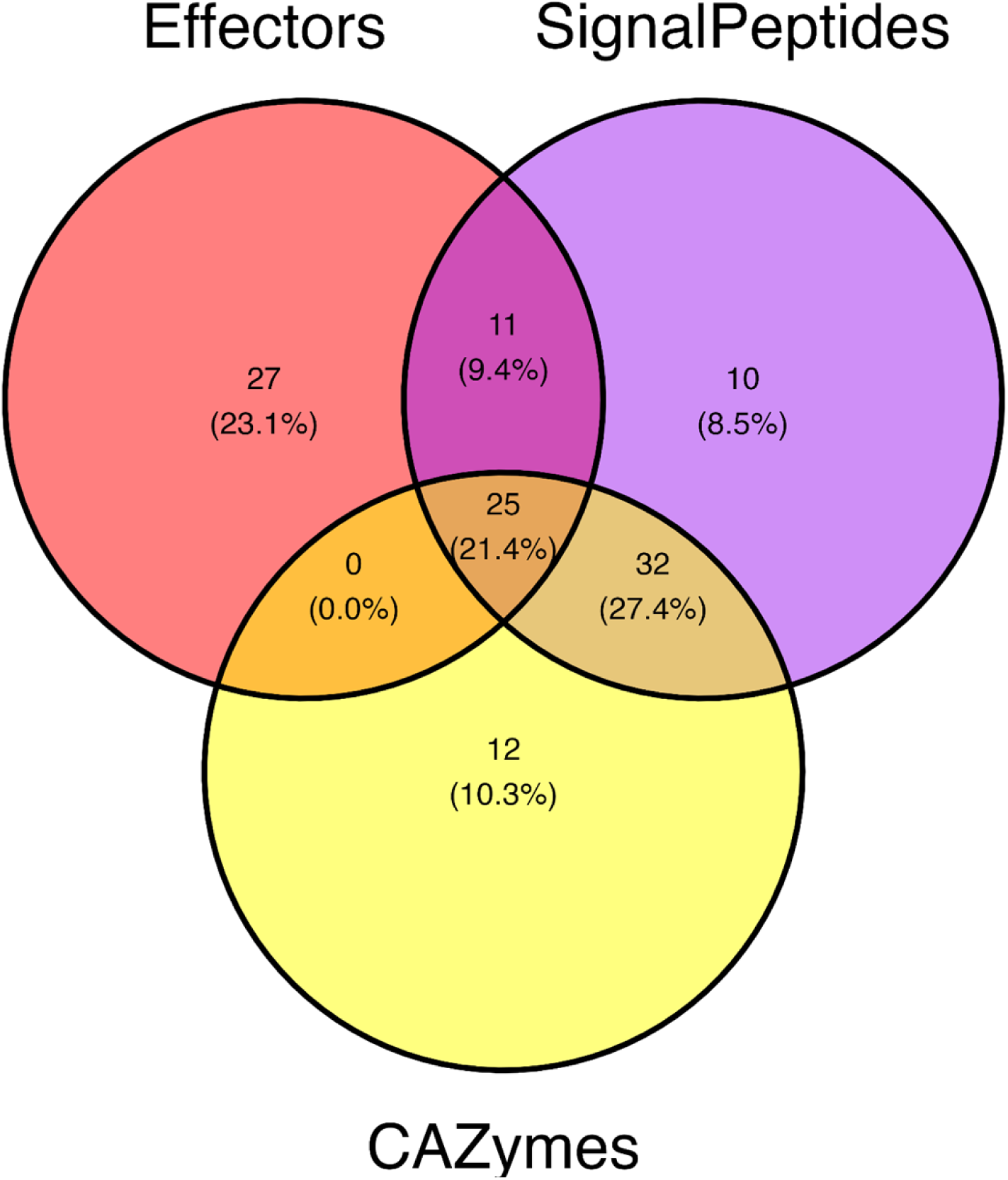
Differentially expressed sporulation genes identified as candidate virulence factors. Venn diagram displaying the number of sporulation genes (*padj* < 0.05) whose predicted protein sequences were also predicted to be at least one candidate virulence factor class: signal peptides, CAZymes (carbohydrate-active enzymes), or effectors. The following were used to identify signal peptides, CAZymes, and effectors: SignapP v5.0, dbCAN v4.1.4, and EffectorP v3.0, respectively.

## Discussion

Here, we present the first T2T, chromosome-level genome assembly and transcriptome study of *D. destructiva* to examine gene expression shifts linked to the transition between vegetative growth and sporulation. This work fills a long-standing gap in genomic resources for both *D. destructiva* and the order Diaporthales more broadly. With this resource, we provide novel insights into the pathogenicity and life cycle regulation of the *D. destructiva* epidemic. A lack of such resources has limited comparative genomics and functional studies across Diaporthales, thereby hindering progress towards understanding the shared evolutionary history, pathogenicity, and management of many of the most devastating forest pathogens. Because *D. destructiva* is strictly anamorphic, conidial sporulation represents the primary means of epidemic-scale dispersal, making it a particularly relevant focus for understanding disease progression and potentially informing strategies for disease management. With this work, we identified key genes and candidate virulence factors, including signal peptides, CAZymes, and effectors, that offer further insight into how *D. destructiva* interacts with *Cornus* species and can facilitate its proliferation within and between susceptible plant hosts. Together, these resources provide foundational tools for comparative and functional genomics and serve as a system for studying other Diaporthalean forest pathogens.

### A comprehensive, annotated genome resource addition to Diaporthales

The newly constructed *D. destructiva* AS111 reference genome generated in this study represents a substantial improvement to resources available for this species and adds a high-quality genomic resource to complement the current resources available for Diaporthales. Six chromosome-level genome assemblies are now available in NCBI that represent various ecologically and economically destructive Diaporthalean phytopathogens [50]. High genome completeness scores from BUSCO, compleasm, and Merqury validate that our assembly accurately represents the *D. destructiva* genome. High genome assembly accuracy is further supported by the high protein completeness scores of the predicted proteome generated from genome annotation. Repeat modeling also highlighted LTR retroelements as the primary repeat elements in *D. destructiva*, which have been implicated in genome evolution and virulence adaptation of other fungal pathogens [81]. Our assembly also includes a circularized mitochondrial genome assembly, which may inform how mitochondria affect metabolic shifts during infection [82–84]. Thus, we present these novel resources as complementary tools for studying fungal genome structure, evolution, and pathogenicity.

### Differential gene expression highlights pathogen competency

Differential expression analysis of sporulating *D. destructiva* (AS111 and WAP31) and *J. japonica* (VA17B) revealed shifts in gene regulation, thereby highlighting key differences in metabolic and functional pathways associated with the induction of sporulation in these closely related fungal taxa. Principal component analysis revealed that interspecific divergence in gene expression exceeded treatment effects, although the latter were still biologically meaningful. The degree of transcriptomics divergence observed between the two *D. destructiva* isolates reflects substantial intra-species variation, consistent with *D. destructiva* isolate divergence previously identified [9]. This variation highlights the importance of incorporating multiple genotypes when investigating transcriptomic responses and functional mechanisms. Among the 162 upregulated genes (*padj* < 0.05), GO enrichment analysis indicated the strongest overrepresentation with hydrolase, polysaccharide, and carbohydrate processes. These enrichments suggest that sporulation genes facilitate cell wall degradation and sugar metabolism to meet structural and energy demands, potentially contributing to the pathogen’s ability to invade and colonize host tissues [85–87]. Four of the most upregulated sporulation genes (*padj* = 1e-7; log2FC = 2) are linked to host tissue degradation, nutrient acquisition, and metabolic adaptation. Xyloglucan endohydrolase (*g8809*) and endo-1,5-α-L-arabinosidase (*g2406*) break down plant cell wall components by targeting xyloglucan and arabinan, respectively [87]. Sugar transport (*g7635*) facilitates nutrient uptake, whereas inositol oxygenase (*g964*) is implicated in metabolic adaptation and virulence [88]. Upregulation of xyloglucan endohydrolase and endo-1,5-α-L-arabinosidase in *D. destructiva* and *J. japonica* highlights the importance of host cell wall degradation during sporulation. Xyloglucan and arabinan are major components of plant cell walls, and their enzymatic breakdown facilitates host tissue penetration [87], thus providing a mechanistic basis for fungal pathogenicity and spore dispersal during the necrotrophic phase. This is particularly crucial for *D. destructiva*, as acervuli must rupture through plant epidermal layers to effectively disperse spores [4].

Inositol oxygenase may also play a crucial role in sporulation by facilitating inositol catabolism, which is vital for fungal development and virulence [88]. In *Cryptococcus neoformans*, inositol metabolism has been linked to the formation of a protective, polysaccharide capsule that imparts pathogenicity by evading immune responses in humans [88]. While true capsule formation is uncommon in phytopathogens, *D. destructiva* has been shown to produce a similar exterior mucilaginous, protein matrix on conidia, which may serve analogous protective functions [39]. The upregulation of inositol oxygenase may contribute to the biosynthesis of these extracellular structures of conidia, potentially influencing spore viability, dispersal, and host attachment during infection. Sugar transport upregulation also suggests an increased demand for nutrient uptake to support the energy-intensive process of sporogenesis. Sugars serve as critical carbon sources, and upregulating transporters may enhance the availability of glucose or other sugars necessary for reproductive metabolic processes [89]. The expression variability observed in *D. destructiva* isolate WAP31, in which sugar transport was downregulated in two biological replicates, may indicate differences in intraspecific sporulation efficiency, but further validation with additional replicates is required to determine whether this reflects technical variation or true biological differences. Similar gene expression patterns of cell wall degradation and nutrient acquisition have been documented in other Diaporthalean phytopathogens, such as *C. parasitica* and *O. clavigignenti-juglandacearum* [90–92]. The conservation of these mechanisms suggests that similar pathogenesis strategies have evolved within Diaporthales to facilitate host invasion and nutrient acquisition. Shared genetic pathways for cell wall degradation and nutrient uptake across Diaporthalean fungi indicate that targeting these mechanisms may be an effective strategy to combat multiple species within this group, including *D. destructiva*.

In contrast, *J. japonica* isolate VA17B exhibited inverse expression patterns for endo-1,5-α-L-arabinosidase, which suggested inter-familial divergence in how these fungi approach polysaccharide degradation during sporulation. However, unique read mapping efficiency among *D. destructiva* isolates WAP31 (western coast genetic cluster) and AS11 (eastern coast genetic cluster) were substantially higher than *J. japonica* isolate VA17B, despite variable intraspecific mapping rates, which may have resulted from regional, intraspecific genetic variation (Table 1) [9]. This divergence in gene expression between the two species underscores the potential for different strategies in tissue degradation and host invasion. Altogether, transcriptional upregulation of these genes suggests that sporulation is not merely a dispersal stage for *D. destructiva*, but also an active process involving host degradation and nutrient acquisition. This finding opens new avenues for understanding the metabolic shifts that support the transition from biotrophy to necrotrophy in *D. destructiva*.

### Metabolic trade-offs during sporulation

Among the 78 downregulated genes (*padj* < 0.05), GO enrichment analysis indicated the strongest overrepresentation with heme binding, tetrapyrrole binding, electron carrier activity, and ion binding metabolic activities. These expression patterns (upregulation and downregulation) suggest that sporulation also involves a reallocation of energy toward reproduction, by prioritizing nutrient acquisition and cell wall degradation to support sporulation while suppressing pathways associated with cellular maintenance and metabolic homeostasis.

The shift in gene regulation indicates that *D. destructiva* and *J. japonica* adjust metabolic strategies to balance the energy demands of reproduction with host tissue degradation.

Four of the most downregulated genes (*padj* = 1e-7; log2FC = 2) contributed to oxidative stress response, nitrogen metabolism, and metal ion regulation. Specifically, ferric-chelate reductase (*g6097*) facilitates iron acquisition; the reactive oxygen species (ROS)-neutralizing gene (*g9869*) detoxifies free radicals; NAD/NADP-dependent octopine/nopaline dehydrogenase (*g5346*) supports nitrogen metabolism; and Ctr copper transporter family genes (*g4351*) regulate copper homeostasis. The concurrent downregulation of copper transporters and iron redox may indicate a broader metabolic reorientation away from metal ion acquisition, in reflection of specific nutrient demands of sporulating cells [93]. This reduction in metal ion acquisition could suggest a fine-tuning of cellular processes to optimize conditions for sporogenesis rather than growth. Similarly, downregulation of ROS-neutralizing genes, such as *g9869*, suggests that *D. destructiva* may exploit oxidative stress as a signaling mechanism in the process of sporulation.

ROS are commonly considered damaging byproducts of metabolism, yet they also play a regulatory role in fungal development, including conidiation and spore germination [94]. Decreased ROS neutralization may highlight a critical aspect of stress adaptation by the pathogen during the reproductive phase, by enabling *D. destructiva* to leverage controlled oxidative stress as a tool for spore development and dispersal, as compared with neutralizing ROS during the vegetative stage. Downregulation of NAD/NADP-dependent octopine/nopaline dehydrogenase (*g5346*), an enzyme involved in nitrogen metabolism, suggests that nitrogen assimilation is less active during sporulation, which is a resource conservation strategy documented in other fungi [95,96].

*Juglanconis japonica* isolate VA17B exhibited a contrasting gene expression pattern under sporulation, with an upregulation of ROS detoxification, nitrogen metabolism, and copper homeostasis genes. This suggests that *J. japonica* employs a different sporulation strategy, possibly requiring tighter regulation of redox balance, energy production, and metal ion homeostasis compared to *D. destructiva*. These differences may reflect species-specific adaptations to the environmental conditions in which they sporulate. For example, *J. japonica* might require more stringent control over ROS and metal ions in its ecological niche, whereas *D. destructiva* may benefit from transient oxidative stress signals that facilitate sporulation.

Although both fungal species share similar ecological niches, their divergent gene expression profiles suggest different molecular strategies for host interaction and reproduction [33].

### Virulence factors reflect hemibiotrophic adaptation

Candidate virulence factor identification revealed enzymatic mechanisms that facilitate host invasion, which underscores the complexity of *D. destructiva*’s hemibiotrophic life cycle. The transition between biotrophic and necrotrophic phases is a hallmark of hemibiotrophic pathogens that requires precise regulation of virulence factors to suppress host defenses early in infection and later, results in degradation of host tissues for nutrient acquisition [35]. The extensive repertoire of CAZymes active in *D. destructiva* suggests key roles for fungal cell wall modification, host tissue degradation, and nutrient acquisition, which are all processes critical for both early colonization and late-stage disease progression [92]. The upregulation of glycoside hydrolases (*g2406* and *g8809*) during sporulation indicates that host cell wall degradation facilitates nutrient uptake required for sporulation. CAZymes are crucial virulence factors in many phytopathogens, with gains or losses of specific CAZyme families often driving evolutionary shifts in pathogenicity [92]. For example, in *C. parasitica*, the loss of specific CAZyme genes has been linked to a transition towards increased pathogenicity, a transition that is absent in related non-pathogenic *Cryphonectria* species [92].

The abundance of predicted signal peptides in *D. destructiva* suggests a rich, diverse secretome that may be critical for both host colonization and tissue necrosis. Signal peptides direct proteins, including effectors, to the secretory pathway, where the proteins are cleaved by signal peptidases and ultimately secreted into the host [78]. Upregulation of the Sec/SPI pathway (*g2406* and *g8809*) during sporulation suggests a complex mechanism for host interactions, potentially contributing to fungal pathogenicity and adaptation. The upregulation of Sec/SPI pathways indicate a sophisticated mechanism for targeting plant cells, in support of the notion that *D. destructiva* relies on both enzymatic degradation and immune suppression to establish infection and promote sustained disease progression in the infected host plant.

Effectors, which manipulate and suppress host immune responses, play a critical role in fungal pathogenicity and are among the most rapidly evolving genes due to frequent mutation and recombination activity [97,98]. These protein effectors are particularly important during the biotrophic phase, when the pathogen must evade recognition and effector-triggered hypersensitivity in the host. The upregulation of *g2406* as a cytoplasmic effector suggests that this gene may facilitate host immune response evasion during sporulation. Contrastingly, downregulation of apoplastic effectors (*g9868*) and cytoplasmic effectors (*g9869*) may indicate a finely tuned regulatory mechanism that tempers aggressive actions of the pathogen during sporulation. However, as infection progresses, it is possible that effector-triggered immunity leading to cell apoptosis could signal a shift from biotrophy to necrotrophy in *D. destructiva*.

### Further time-course research is needed to validate this relationship

Altogether, our study provides insight into virulence mechanisms and metabolic shifts that support infection and sporulation in *D. destructiva*. These findings contextualize the gene expression profile of *D. destructiva* at critical life cycle transitions, as well as offer novel insight into potential virulence mechanisms in one of the most severe forest epidemics of the 20th century in North America. This work also strengthens the foundation for comparative genomics with other Diaporthalean pathogens, which are known or are positioned to negatively impact forest ecosystems. Understanding genome-scale diversity among pathogens will enhance our ability to forecast future biosecurity risks. As anthropogenic factors such as global trade and climate change continue to facilitate the spread of pathogens, integrating such multi-omics resources into early detection frameworks and risk assessment models will be essential for proactive conservation and the safeguarding of forest ecosystems.

## Acknowledgements

We gratefully recognize the UTIA Genomics Center for the Advancement of Agriculture and the UT Genomics Core for laboratory supplies and conducting RNA sequencing. We also thank the University of Washington Northwest Genomics Center for PacBio HiFi sequencing and Phase Genomics for Hi-C sequencing. Thank you to Mark Windham (University of Tennessee) and Ning Zhang (Rutgers University) for sharing historical fungal isolates that were vital to this research. Special thanks to the following people for their helpful feedback and suggestions throughout this research: Denita Hadziabdic-Guerry, DeWayne D. Shoemaker, Sarah L. Boggess, Lav K. Yadav, Ryan Kuster, Aaron Onufrak, Trinity P. Hamm, Zane C. Smith, Alina Pokhrel, and Kayleigh P. Redington.

## Data Availability

The raw data and annotated genome assembly for this study have been deposited in the European Nucleotide Archive (ENA) at EMBL-EBI under the study accession number PRJEB90672 (https://www.ebi.ac.uk/ena/browser/view/PRJEB90672). Code for all genome assembly, annotation, and RNAseq analyses can be found in the following GitHub wikis: https://github.com/ShadeNiece/DisculaDestructiva_GenomeAssembly-Annotation/wiki and https://github.com/ShadeNiece/DisculaDestructiva_RNAseq/wiki, respectively.

## Supporting Information

**Fig S1. Images of RNAseq cultures on sporulating and non-sporulating media.** Images on the left represent non-sporulating cultures plated on half strength PDA^++^. Images to the right represent sporulating cultures grown on solid water agar supplemented with finely ground *Quercus alba* L. (white oak) leaf tissue. A magnified image is included for each to show the presence of strictly mycelia (left) and both mycelia and acervuli (right). The isolate shown is *Discula destructiva* isolate AS111.

**Fig S2. Telomeric signal plot for the nuclear genome assembly of *Discula destructiva*.** Telomeric signal plot from tidk v0.2.65 showing the distribution of telomeric motifs 5’-(TTAGGG/CCCTAA)_n_-3 across the eight chromosome-level scaffolds in the *D. destructiva* AS111 nuclear genome assembly. Peaks at the beginning and end of each scaffold indicate the presence of telomeric repeats. Scaffold lengths are shown in megabases (Mb).

**Fig S3. PCA plot of gene expression variance among *Discula destructiva* isolates (AS111 and WAP31) and *Juglanconis japonica* (isolate VA17B).** The principal component analysis (PCA) was generated using DESeq2 (v1.42.1). The x-axis (PC1) and y-axis (PC2) explain 54% and 16% of the total variance, respectively. Each point represents a sample (n = 18), color-coded by growth medium: sporulating treatment (OAK–red) and non-sporulating treatment (PDA–teal). Black ellipses highlight sample clustering within each isolate. Separation along PC1 reflects inter-familial divergence between *D. destructiva* and *J. japonica*, while PC2 reflects intra-species variation within *D. destructiva*. AS111 belongs to the southern region genetic cluster (collected in 2000), and WAP31 to the western region cluster (collected in 1989) (Mantooth et al., 2017).

**Table S1. Diaporthalean chromosome-scale genome assemblies on NCBI.** Current chromosome-level genome assemblies for five economically and ecologically destructive Diaporthalean pathogens available in NCBI.

**Table S2. List of Abbreviations.** List of commonly used abbreviations throughout this manuscript.

**Table S3. Raw HiFi long-read sequencing statistics for *Discula destructiva* isolate AS111.**

Statistics were provided from the raw Pacbio HiFi run using the Revio system.

**Table S4. Genome assembly statistics for the nuclear genome assembly of *Discula destructiva* isolate AS111.** Genome statistics generated using the BBMap v39.06 stats function. Contig-level assembly refers to the assembly prior to scaffolding, mitochondrion filtering, gap closing, and telomere-to-telomere (T2T) verification. Chromosome-level assembly refers to the assembly after scaffolding and mitochondrion filtering, but before gap closing and T2T verification. Final T2T assembly refers to the assembly after scaffolding, mitochondrial filtering, gap closing, and T2T verification.

**Table S5. Genome annotation statistics for the nuclear genome assembly of *Discula destructiva* isolate AS111.** Structural annotations were performed using BRAKER v3.0.8. Repeats were modeled using RepeatModeler v2.0.5 and masked using RepeatMasker v4.1.6. These statistics were generated by mapping all raw RNAseq reads for *D. destructiva* (isolates AS111 and WAP31) and *J. japonica* (isolate VA17B) to the *D. destructiva* AS111 reference genome.

**Table S6. RNAseq read statistics for *Discula destructiva* and *Juglanconis japonica*.** RNA sequencing was performed with a SP, 300 cycle flow cell on a NovaSeq platform. Statistics for pre- and post-trimmed RNAseq read-pairs were generated using FastQC v0.11.7. Quality trimming of RNAseq read-pairs was performed with BBDuk v39.06.

**Table S7. Mapping rates of *Discula destructiva* and *Juglanconis japonica* RNAseq read-pairs.** Reads were mapped to the *D. destructiva* AS111 reference genome using STAR v2.7.11b. OAK: sporulating treatment (mycelia and acervuli present). PDA: non-sporulating treatment (strictly mycelia).

**Table S8. 240 differentially expressed genes identified under the sporulation treatment.**

Significance was identified using an *adjusted P value* (*padj*) < 0.05 via DESeq2 v1.42.1. Of the 240 genes, 162 genes were upregulated, and 78 genes were downregulated. Reads from *D. destructiva* (isolates AS111 and WAP31) and *J. japonica* (isolate VA17-B) were mapped to the *D. destructiva* AS111 reference genome using STAR v2.7.11b.

**Table S9. KEGG Ontology (KO) annotations for 240 significant sporulation genes.** Significance was identified using an *adjusted P value* (*padj*) < 0.05 via DESeq2 v1.42.1. Multiple KO annotations may be displayed for one gene. “NA” represents genes which could not be functionally annotated with EnTAP v1.0.0.

**Table S10. Candidate virulence factors identified for *Discula destructiv*a isolate AS111.** These virulence factors were predicted using the *D. destructiva* isolate AS111 proteome that was generated by BRAKER v3.0.8. A total of 11,480 protein sequences were profiled for each of the three virulence factors. Signal peptides were predicted using SignalP v5.0. CAZymes were predicted using dbCAN v4.1.4. Effectors were predicted using EffectorP v3.0.

**Table S11. Gene IDs and their predictions as signal peptides.** SignalP v5.0 was used to predict signal peptides from 11,480 protein sequences using the *Discula destructiva* AS111 proteome.

**Table S12. Gene IDs and their predictions as carbohydrate-active enzymes (CAZymes).** DbCAN v4.1.4 was used to predict CAZymes from 11,480 protein sequences using the *Discula destructiva* AS111 proteome.

**Table S13. Gene IDs and their predictions as effectors.** EffectorP v3.0 was used to predict effectors from 11,480 protein sequences using the *Discula destructiva* AS111 proteome.

**Table S14. Gene IDs of differentially expressed sporulation genes identified as candidate virulence factors.** Of the 240 differentially expressed sporulation genes (padj < 0.05), 117 were identified to have predicted protein sequences that overlapped with at least one of the three virulence factors (e.g. signal peptides, CAZymes, or effectors). Signal peptides were predicted using SignalP v5.0. CAZymes were predicted using dbCAN v4.1.4. Effectors were predicted using EffectorP v3.0.33

**File S1. HMW DNA Extraction Protocol for *Discula destructiva*.** HMW DNA extraction protocol optimized for the filamentous fungus, *Discula destructiva*.

**File S2. Polysaccharide Purification Protocol.** High salt, low ethanol polysaccharide precipitation to purify genomic DNA for PacBio HiFi sequencing.

**File S3. RNA Extraction Protocol.** RNA extraction protocol optimized for the filamentous fungi, *Discula destructiva* and *Juglanconis japonica*.

**File S4. Mt_removal.py.** Python script to identify mitochondrial sequences in the nuclear assembly by finding and removing any contigs with aligned regions that have more than 90% alignment.

**File S5. DGE_Analysis.R.** Differential gene expression analysis of sporulating samples using DESeq2.

**File S6. GO_Enrichment.R.** GO and KO Associations for differentially expressed sporulation genes.

**File S7. Virulence_Factors.R.** Overlap Between Virulence Factors and Sporulation Genes.

## References

1. Castlebury LA, Rossman AY, Jaklitsch WJ, Vasilyeva LN. A preliminary overview of the Diaporthales based on large subunit nuclear ribosomal DNA sequences. Mycologia. 2002;94(6):1017–31.

2. Sogonov MV, Castlebury LA, Rossman AY, Mejía LC, White JF. Leaf-inhabiting genera of the Gnomoniaceae, Diaporthales. Stud Mycol. 2008;62:1–77.

3. Senanayake IC, Crous PW, Groenewald JZ, Maharachchikumbura SSN, Jeewon R, Phillips AJL, et al. Families of Diaporthales based on morphological and phylogenetic evidence. Stud Mycol. 2017;86:217–96.

4. Redlin SC. *Discula destructiva* sp. nov., cause of dogwood anthracnose. Mycologia. 1991;83(5):633–42.

5. Potter KM, Escanferla ME, Jetton RM, Man G, Crane BS. Prioritizing the conservation needs of United States tree species: Evaluating vulnerability to forest insect and disease threats. Glob Ecol Conserv. 2019;18:e00622.

6. Hadziabdic D, Fitzpatrick BM, Wang X, Wadl PA, Rinehart TA, Ownley BH, et al. Analysis of genetic diversity in flowering dogwood natural stands using microsatellites: The effects of dogwood anthracnose. Genetica. 2010;138(9):1047–57.

7. Oswalt CM, Oswalt SN, Woodall CW. An assessment of flowering dogwood (*Cornus florida* L.) decline in the Eastern United States. Open J For. 2012;02(02):41.

8. Eyde RH. Comprehending *Cornus*: Puzzles and progress in the systematics of the dogwoods. Bot Rev. 1988;54(3):233–351.

9. Mantooth K, Hadziabdic D, Boggess SL, Windham MT, Miller S, Cai G, et al. Confirmation of independent introductions of an exotic plant pathogen of *Cornus* species, *Discula destructiva*, on the east and west coasts of North America. PLOS ONE. 2017;12(7):e0180345.

10. Mielke M, Langdon K. Dogwood anthracnose fungus threatens Catoctin Mt. Park. Park Sci. 1986;6(2):6–8.

11. Caetano-Anollés G, Trigiano RN, Windham MT. Patterns of evolution in *Discula* fungi and the origin of dogwood anthracnose in North America, studied using arbitrarily amplified and ribosomal DNA. Curr Genet. 2001;39(5):346–54.

12. Orwig DA, Abrams MD. Land-use history (1720–1992), composition, and dynamics of oak–pine forests within the Piedmont and Coastal Plain of northern Virginia. Can J For Res. 1994;24(6):1216–25.

13. Jenkins MA, Parker GR. Composition and diversity of woody vegetation in silvicultural openings of southern Indiana forests. For Ecol Manag. 1998;109(1):57–74.

14. Jenkins MA, Jose S, White PS. Impacts of an exotic disease and vegetation change on foliar calcium cycling in Appalachian forests. Ecol Appl. 2007;17(3):869–81.

15. Holzmueller EJ, Jose S, Jenkins MA. Relationship between *Cornus florida* L. and calcium mineralization in two southern Appalachian forest types. For Ecol Manag. 2007;245(1):110–7.

16. Pollock MM, Beechie TJ, Imaki H. Using reference conditions in ecosystem restoration: An example for riparian conifer forests in the Pacific Northwest. Ecosphere. 2012;3(11):art98.

17. Dirr MA. Manual of woody landscape plants: Their identification, ornamental characteristics, culture, propagation and uses. 6th Edition. Stipes Publishing, Champaign, IL; 2009.

18. Nowicki M, Houston LC, Boggess SL, Aiello AS, Payá-Milans M, Staton ME, et al. Species diversity and phylogeography of *Cornus kousa* (Asian dogwood) captured by genomic and genic microsatellites. Ecol Evol. 2020;10(15):8299–312.

19. Census of Horticulture Specialties. Washington, D.C.: U.S. Department of Agriculture National Agriculture Statistics Service; 2020.

20. Moreau ELP, Medberry AN, Honig JA, Molnar TJ. Genetic diversity analysis of big-bracted dogwood (*Cornus florida* and *C. kousa*) cultivars, interspecific hybrids, and wild-collected accessions using RADseq. PLOS ONE. 2024;19(7):e0307326.

21. Nowicki M, Boggess SL, Saxton AM, Hadziabdic D, Xiang QYJ, Molnar TJ, et al. Haplotyping of Cornus florida and C. kousa chloroplasts: Insights into species-level differences and patterns of plastic DNA variation in cultivars. Heinze B, editor. PLOS ONE. 2018;13(10):e0205407.

22. Windham MT, Graham ET, Witte WT, Knighten JL, Trigiano RN. *Cornus florida* ‘Appalachian Spring’: A white flowering dogwood resistant to dogwood anthracnose. HortScience. 1998;33(7):1265–7.

23. Daughtrey ML, Hibben CR. Lower branch dieback, a new disease of northern dogwoods. Phytopathology. 1983;73(365).

24. Hibben CR, Daughtrey ML. Dogwood anthracnose in Northeastern United States. Plant Dis. 1988;72(3):199.

25. Anderson RL, Knighten JL, Windham MT, Langdon K, Hedrix F, Roncadori R. Dogwood anthracnose and its spread in the South. U.S. Department of Agriculture Forest Service Southern Region. 1994.

27. Byther RS, Davidson Jr. RM. Dogwood anthracnose. Coop Ext Serv. 1979;3:20–1.

28. Pirone PP. Parasitic fungus affects region’s dogwood; Dogwood disease [Internet]. The New York Times. 1980 [cited 2025 Jan 22]. Available from: https://www.nytimes.com/1980/02/24/archives/parasitic-fungus-affects-regions-dogwood-dogwood-disease.html

29. Voglmayr H, Castlebury LA, Jaklitsch WM. *Juglanconis* gen. nov. on Juglandaceae, and the new family Juglanconidaceae (Diaporthales). Persoonia - Mol Phylogeny Evol Fungi. 2017;38(1):136–55.

30. Ownley BH, Trigiano RN. Plant pathology concepts and laboratory exercises. 3rd Edition. Taylor & Francis; 2017.

31. Davidson RW, Campbell WA, Blaisdell DJ. Differentiation of wood-decaying fungi by their reactions on gallic or tannic acid medium. J Agric Res. 1938;57:683–95.

32. Trigiano RN, Gerhaty NE, Windham MT, Brown DA. A simple assay for separating fungi associated with dogwood anthracnose. In: Proceedings of Southern Nursery Association Research Conference. 1991. p. 209–11.

33. Nowicki M, Redington KP, Boggess SL, Niece IS, and Trigiano RN. The enzymatic arsenal of *Discula destructiva*: Strategies for understanding and managing the dogwood anthracnose pathogen. Mycologia. 0(0):1–14.

34. Smith-Salogga D. Occurrence, symptoms and probable cause, Discula sp., of Cornus leaf anthracnose. [Master’s Thesis]. [Seattle, WA]: University of Washington, Seattle; 1982.

35. Perfect SE, Green JR. Infection structures of biotrophic and hemibiotrophic fungal plant pathogens. Mol Plant Pathol. 2001;2(2):101–8.

36. Wedge DE, Riley MB, Tainter FH. Phytotoxicity of *Discula destructiva* culture filtrates to *Cornus* spp. and the relationship to disease symptomology. Plant Dis. 1999;83(4):377–80.

37. Venkatasubbaiah P, Chilton WS. Toxins produced by the dogwood anthracnose fungus *Discula* sp. J Nat Prod. 1991;54(5):1293–7.

38. Daughtrey ML, Hibben CR, Britton KO, Windham MT, Redlin SC. Dogwood anthracnose: Understanding a disease new to North America. Plant Dis. 1996;80(4):349.

39. Colby DM, Windham MT, Grant JF. Transportation and viability of conidia of *Discula destructiva* on *Hippodamia convergens*. Plant Dis. 1996;80(7):804.

40. Cornus florida (Flowering Dogwood) [Internet]. NatureServe Explorer. [cited 2024 July 28]. Available from: https://explorer.natureserve.org/Taxon/ELEMENT_GLOBAL.2.149695/Cornus_florida

41. Mitchell E, Fleming S, Dorken M, Freeland J. Susceptibility of endangered *Cornus florida* (eastern flowering dogwood) to the introduced fungal pathogen *Discula destructiva* (dogwood anthracnose) in the Canadian Carolinian forest: Insights from environmental, ecological, and population genetics assessments. Botany. 2023;cjb-2022-0088.

42. Cornus nuttallii (Pacific Dogwood) [Internet]. NatureServe Explorer. [cited 2024 July 28]. Available from: https://explorer.natureserve.org/Taxon/ELEMENT_GLOBAL.2.159922/Cornus_nuttallii

43. Trigiano RN, Hamm TP, Boggess SL, Staton ME. ‘Rebecca’s Appalachian Angel’: A cultivar of flowering dogwood (*Cornus florida*) with large leaves and floppy white bracts. HortScience. 2023;58(8):881–4.

44. Fulcher A, Hale F, Windham A. Dogwood—Cornus spp. IPM for select deciduous trees in Southeastern US nursery production. South Nurs IPM Work Group. 2012.

45. Santamour, Jr. FS, McArdle AJ, Strider PV. Susceptibility of flowering dogwood of various provenances to dogwood anthracnose. Plant Dis. 1989;73(7):590.

46. Hadziabdic D, Hamelin RC, Stewart JE, Villari C. Editorial: Forest pathology in changing climate. Front For Glob Change [Internet]. 2022 Sept 30 [cited 2024 Nov 1];5. Available from: https://www.frontiersin.org/journals/forests-and-global-change/articles/10.3389/ffgc.2022.1032035/full

47. Miller S, Masuya H, Zhang J, Walsh E, Zhang N. Real-time PCR detection of dogwood anthracnose fungus in historical herbarium specimens from Asia. PLoS ONE. 2016;11(4):e0154030.

48. Trigiano RN, Caetano-Anollés G, Bassam BJ, Windham MT. DNA amplification fingerprinting provides evidence that *Discula destructiva*, the cause of dogwood anthracnose in North America, is an introduced pathogen. Mycologia. 1995;87(4):490–500.

49. Zhang N, Blackwell M. Population structure of dogwood anthracnose fungus. Phytopathology®. 2002;92(12):1276–83.

50. Sayers EW, Bolton EE, Brister JR, Canese K, Chan J, Comeau DC, et al. Database resources of the national center for biotechnology information. Nucleic Acids Res. 2022;50(D1):D20–6.

51. McElreath SD, Tainter FH. A sporulation medium for *Discula destructiva*, the dogwood anthracnose fungus. Curr Microbiol. 1993;26(2):117–21.

52. Andrews S. FastQC: A quality control tool for high throughput sequence data [Internet]. Babraham Bioinformatics. [cited 2025 Jan 26]. Available from: https://www.bioinformatics.babraham.ac.uk/projects/fastqc/

53. Seqtk: Toolkit for processing sequences in FASTA/Q formats [Internet]. lh3/seqtk. [cited 2024 July 7]. Available from: https://github.com/lh3/seqtk

54. Cheng H, Concepcion GT, Feng X, Zhang H, Li H. Haplotype-resolved *de novo* assembly using phased assembly graphs with hifiasm. Nat Methods. 2021;18(2):170–5.

55. Astashyn A, Tvedte ES, Sweeney D, Sapojnikov V, Bouk N, Joukov V, et al. Rapid and sensitive detection of genome contamination at scale with FCS-GX. Genome Biol. 2024;25(1):60.

56. Zhou C, Brown M, Blaxter M, Consortium TDT of LP, McCarthy SA, Durbin R. Oatk: A de novo assembly tool for complex plant organelle genomes [Internet]. bioRxiv; 2024 [cited 2025 Feb 12]. p. 2024.10.23.619857. Available from: https://www.biorxiv.org/content/10.1101/2024.10.23.619857v1

57. Bushnell B. BBMap: A fast, accurate, splice-aware aligner. Berkeley Natl Lab [Internet]. 2014 Mar 19 [cited 2025 Feb 12];(LBNL Report #: LBNL-7065E). Available from: https://escholarship.org/uc/item/1h3515gn

58. Tillich M, Lehwark P, Pellizzer T, Ulbricht-Jones ES, Fischer A, Bock R, et al. GeSeq – versatile and accurate annotation of organelle genomes. Nucleic Acids Res. 2017;45(W1):W6–11.

59. Greiner S, Lehwark P, Bock R. OrganellarGenomeDRAW (OGDRAW) version 1.3.1: Expanded toolkit for the graphical visualization of organellar genomes. Nucleic Acids Res. 2019;47(W1):W59–64.

60. Zhou C, McCarthy SA, Durbin R. YaHS: Yet another Hi-C scaffolding tool. Alkan C, editor. Bioinformatics. 2023;39(1):btac808.

61. Durand NC, Robinson JT, Shamim MS, Machol I, Mesirov JP, Lander ES, et al. Juicebox provides a visualization system for Hi-C contact maps with unlimited zoom. Cell Syst. 2016;3(1):99–101.

62. Xu M, Guo L, Gu S, Wang O, Zhang R, Peters BA, et al. TGS-GapCloser: A fast and accurate gap closer for large genomes with low coverage of error-prone long reads. GigaScience. 2020;9(9):giaa094.

63. Brown MR, Manuel Gonzalez de La RosaP, Blaxter M. Tidk: A toolkit to rapidly identify telomeric repeats from genomic datasets. Bioinformatics. 2025;41(2):btaf049.

64. Simão FA, Waterhouse RM, Ioannidis P, Kriventseva EV, Zdobnov EM. BUSCO: Assessing genome assembly and annotation completeness with single-copy orthologs. Bioinformatics. 2015;31(19):3210–2.

65. Huang N, Li H. Compleasm: A faster and more accurate reimplementation of BUSCO. Bioinformatics. 2023;39(10):btad595.

66. Rhie A, Walenz BP, Koren S, Phillippy AM. Merqury: Reference-free quality, completeness, and phasing assessment for genome assemblies. Genome Biol. 2020;21(1):245.

67. Flynn JM, Hubley R, Goubert C, Rosen J, Clark AG, Feschotte C, et al. RepeatModeler2 for automated genomic discovery of transposable element families. Proc Natl Acad Sci. 2020;117(17):9451–7.

68. Smit A, Hubley R, Green P. RepeatMasker Open-4.0 [Internet]. 2013. Available from: http://www.repeatmasker.org

69. Dobin A, Davis CA, Schlesinger F, Drenkow J, Zaleski C, Jha S, et al. STAR: Ultrafast universal RNA-seq aligner. Bioinformatics. 2013;29(1):15–21.

70. Gabriel L, Brůna T, Hoff KJ, Ebel M, Lomsadze A, Borodovsky M, et al. BRAKER3: Fully automated genome annotation using RNA-seq and protein evidence with GeneMark-ETP, AUGUSTUS, and TSEBRA. Genome Res [Internet]. 2024 June 12 [cited 2025 Feb 12]; Available from: https://genome.cshlp.org/content/early/2024/05/28/gr.278090.123

71. Hart AJ, Ginzburg S, Xu M (Sam), Fisher CR, Rahmatpour N, Mitton JB, et al. EnTAP: Bringing faster and smarter functional annotation to non-model eukaryotic transcriptomes. Mol Ecol Resour. 2020;20(2):591–604.

72. Liao Y, Smyth GK, Shi W. FeatureCounts: An efficient general purpose program for assigning sequence reads to genomic features. Bioinformatics. 2014;30(7):923–30.

73. Love MI, Huber W, Anders S. Moderated estimation of fold change and dispersion for RNA-seq data with DESeq2. Genome Biol. 2014;15(12):550.

74. Posit Team. RStudio: Integrated Development Environment for R [Internet]. Boston, MA: Posit Studio, PBC; 2024. Available from: http://www.posit.co/

75. Ogata H, Goto S, Sato K, Fujibuchi W, Bono H, Kanehisa M. KEGG: Kyoto Encyclopedia of Genes and Genomes. Nucleic Acids Res. 1999;27(1):29–34.

76. Kanehisa M, Sato Y, Morishima K. BlastKOALA and GhostKOALA: KEGG tools for functional characterization of genome and metagenome sequences. J Mol Biol. 2016;428(4):726–31.

77. Jones P, Binns D, Chang HY, Fraser M, Li W, McAnulla C, et al. InterProScan 5: Genome-scale protein function classification. Bioinformatics. 2014;30(9):1236–40.

78. Almagro Armenteros JJ, Tsirigos KD, Sønderby CK, Petersen TN, Winther O, Brunak S, et al. SignalP 5.0 improves signal peptide predictions using deep neural networks. Nat Biotechnol. 2019;37(4):420–3.

79. Zheng J, Ge Q, Yan Y, Zhang X, Huang L, Yin Y. DbCAN3: Automated carbohydrate-active enzyme and substrate annotation. Nucleic Acids Res. 2023;51(W1):W115–21.

80. Sperschneider J, Dodds PN. EffectorP 3.0: Prediction of apoplastic and cytoplasmic effectors in fungi and oomycetes. Mol Plant-Microbe Interactions®. 2022;35(2):146–56.

81. Castanera R, López-Varas L, Borgognone A, LaButti K, Lapidus A, Schmutz J, et al. Transposable elements versus the fungal genome: Impact on whole-genome architecture and transcriptional profiles. PLOS Genet. 2016;12(6):e1006108.

82. Yildiz G, Ozkilinc H. First characterization of the complete mitochondrial genome of fungal plant-pathogen *Monilinia laxa* which represents the mobile intron rich structure. Sci Rep. 2020;10(1):13644.

83. Mendoza H, Perlin MH, Schirawski J. Mitochondrial inheritance in phytopathogenic fungi—everything is known, or is it? Int J Mol Sci. 2020;21(11):3883.

84. Baidyaroy D, Huber DH, Fulbright DW, Bertrand H. Transmissible mitochondrial hypovirulence in a natural population of *Cryphonectria parasitica*. Mol Plant-Microbe Interactions®. 2000;13(1):88–95.

85. Stajich JE. Fungal genomes and insights into the evolution of the kingdom. Microbiol Spectr. 2017;5(4):10.1128/microbiolspec.FUNK-0055–2016.

86. Sun P, Li X, Dilokpimol A, Henrissat B, de Vries RP, Kabel MA, et al. Fungal glycoside hydrolase family 44 xyloglucanases are restricted to the phylum Basidiomycota and show a distinct xyloglucan cleavage pattern. iScience. 2021;25(1):103666.

87. Ransom RF, Walton JD. Purification and characterization of extracellular β-xylosidase and α-arabinosidase from the plant pathogenic fungus *Cochliobolus carbonum*. Carbohydr Res. 1997;297(4):357–64.

88. Wang Y, Wear M, Kohli G, Vij R, Giamberardino C, Shah A, et al. Inositol metabolism regulates capsule structure and virulence in the human pathogen *Cryptococcus neoformans*. mBio. 2021;12(6):e02790–21.

89. Zheng J, Hu B, Zhang X, Ge Q, Yan Y, Akresi J, et al. DbCAN-seq update: CAZyme gene clusters and substrates in microbiomes. Nucleic Acids Res. 2023;51(D1):D557–63.

90. Kubicek CP, Starr TL, Glass NL. Plant cell wall–degrading enzymes and their secretion in plant-pathogenic fungi. Annu Rev Phytopathol. 2014;52(Volume 52, 2014):427–51.

91. Wu G, Schuelke TA, Iriarte G, Broders K. The genome of the butternut canker pathogen, *Ophiognomonia clavigignenti-juglandacearum* shows an elevated number of genes associated with secondary metabolism and protection from host resistance responses. PeerJ. 2020;8:e9265.

92. Stauber L, Prospero S, Croll D. Comparative genomics analyses of lifestyle transitions at the origin of an invasive fungal pathogen in the genus *Cryphonectria*. mSphere. 2020;5(5):e00737–20.

93. Robinson JR, Isikhuemhen OS, Anike FN. Fungal–metal interactions: A review of toxicity and homeostasis. J Fungi. 2021 18;7(3):225.

94. Heller J, Tudzynski P. Reactive oxygen species in phytopathogenic fungi: Signaling, development, and disease. Annu Rev Phytopathol. 2011;49(Volume 49, 2011):369–90.

95. Lee IR, Chow EWL, Morrow CA, Djordjevic JT, Fraser JA. Nitrogen metabolite repression of metabolism and virulence in the human fungal pathogen *Cryptococcus neoformans*. Genetics. 2011;188(2):309–23.

96. Donofrio NM, Oh Y, Lundy R, Pan H, Brown DE, Jeong JS, et al. Global gene expression during nitrogen starvation in the rice blast fungus, *Magnaporthe grisea*. Fungal Genet Biol. 2006;43(9):605–17.

97. Lo Presti L, Lanver D, Schweizer G, Tanaka S, Liang L, Tollot M, et al. Fungal effectors and plant susceptibility. Annu Rev Plant Biol. 2015;66(1):513–45.

98. Möller M, Stukenbrock EH. Evolution and genome architecture in fungal plant pathogens. Nat Rev Microbiol. 2017;15(12):756–71.

